# Inhibitory Plasticity Enhances Sequence Storage Capacity and Retrieval Robustness

**DOI:** 10.1101/2024.04.08.588573

**Authors:** Ziyi Gong, Nicolas Brunel

**Affiliations:** Department of Neurobiology, Duke University, Durham, NC, USA; Department of Physics, Duke University, Durham, NC, USA; Department of Computing Sciences, Bocconi University, Milan, Italy

## Abstract

The generation of motor behaviors and the performance of complex cognitive tasks rely on sequential activity in specific brain structures. The mechanisms of learning and retrieval of these temporal patterns of activity are still poorly understood. Emerging evidence has highlighted the importance of inhibition to learning and memory. However, the specific functions of inhibitory plasticity in the learning and retrieval of sequential activity have been studied very little, apart from its role in maintaining excitation-inhibition (E-I) balance. Using simulations and dynamical mean-field theory of balanced E-I networks, we found that sequences can be stored and retrieved using plasticity in both E-to-I and I-to-E pathways, in the absence of recurrent excitatory plasticity. Networks with both E-to-I and I-to-E plasticity are shown to exhibit higher optimal capacity than models in which plasticity is restricted to recurrent excitation. We further show that inhibitory plasticity enhances robustness to external noise and initial cue perturbation. Thus, our work suggests new computational roles for inhibitory plasticity in improving capacity and robustness of sequence learning.

## 1 Introduction

Animals interact with temporally varying environments. It is crucial for the brain to support a variety of cognitive processing and generate diverse behaviors over time. Experiments in various animals suggest that complex behaviors and cognitive functions rely on sequential activity in specific brain structures. For example, sTable sequential firing of pyramidal cells in motor cortex emerges when mice learn to produce specific movements [1, 39]. Similar spatiotemporal patterns of activity exist in other brain structures during various types of behavior. In zebra finches, HVC neurons burst sequentially during song production [17], and perturbing this sequential firing disrupt the structure of song syllables[55]. Beside motor functions, sequential activity has also been suggested to encode time [20, 42, 44, 56], spatial trajectories [28], and choice-related information [19]. Therefore, sequential activity may play important roles in multiple functions. However, how these temporal patterns are stored and robustly generated are not well understood.

To explore the mechanisms of sequence storage and generation, numerous theoretical models have been proposed. A widely studied class of sequence-generating models is the “synfire-chain” [5, 11, 13], where clusters of excitatory neurons are connected into one or multiple chains. Synchronous firing of neurons in one cluster excites its downstream clusters with a delay, creating sequences of active clusters. The synfire-chain model has received support from recordings of HVC, a structure in songbirds that generates precise sequential bursting activity and is critical for vocal production [31]. However, this model fails to capture observed characteristics of sequential activity in other cortical structures in which neurons show diverse activity profiles with broader temporal activation[1, 19, 44, 47, 56].

Another class of models involves training recurrent neural networks (RNNs) via supervised learning to generate specific spatiotemporal patterns [18, 27, 46]. The produced sequential activity can match experimental observations well, and can be robust to noise, compared to a chaotic, randomly connected RNN [27]. However, these models rely on heavily supervised teaching signals to stably generate the sequences. It is unclear whether biological circuits can implement the non-local learning rules used to update the synaptic weights in such networks.

Several studies have shown that RNNs can also store and retrieve sequences with more biologically plausible learning rules, such as temporally asymmetric Hebbian (TAH) learning rules [15, 16, 24, 25, 38]. TAH rules strengthen (weaken) a synapse if the post-synaptic neuron is activated at a certain delay after (before) the activation of the pre-synaptic neuron. A recent study [15] shows that RNNs endowed with a TAH rule can store sequences and retrieve them with characteristics matching experimental observations. For instance, neurons at the beginning of the sequential activity are more sharply tuned compared to those toward the end, and more neurons code for the initial phase of a retrieved sequence, compared to later phases[1, 44, 47].

All models described so far either do not respect Dale’s law, or involve only recurrent excitatory (E→ E) plasticity. However, emerging evidence suggests that inhibitory neurons are involved in different forms of synaptic plasticity [4, 9, 14, 22, 23, 26, 32, 49, 53, 57], and may play important roles in sequence learning [1, 4, 22, 29, 49], in addition to maintaining E-I balance. For example, in spatial navigation tasks, up- and down-regulating LTP at excitatory synapses onto hippocampal CA1 somatostasin-expressing interneurons (SOM-INs) bidirectionally regulate spatial memory [4]. During motor learning, the axonal boutons of SOM-INs and parvalbumin-expressing inhibitory neurons (PV-INs) in the primary motor cortex (M1) decreased and increased respectively [9]. Manipulating SOM-IN activity disrupted the stability of dendritic spines of excitatory neurons and impaired the establishment of stereotyped-movements. In another study, activating SOM-INs during motor learning de-stabilized learning-induced sequential activity in M1 [1]. Together, these results suggest critical functions of inhibitory plasticity in learning that need to be explored.

To investigate the roles of inhibitory plasticity, we built balanced excitation-inhibition (E-I) networks with different combinations of Hebbian plasticity rules. We found that networks with plasticity in both E→ I and I→ E pathways can store and retrieve sequences, in the absence of E→ E plasticity. We estimated the storage capacity of sequences in different models as a function of learning-related parameters. A grid search over these parameters showed that the networks storing sequences without E→ E plasticity had the largest optimal capacity, in terms of number of patterns stored per plastic synapse. During retrieval of sequences, inhibitory plasticity also enhanced robustness to external noise and initial cue perturbation. In summary, our work suggests new computational roles for inhibitory plasticity in improving capacity and robustness of sequence learning.

## 2 Results

### 2.1 E-I Network Model

To investigate the impact of inhibitory plasticity on sequence learning and retrieval, we analyzed a sparsely connected E-I network consisting of *N*_*E*_ excitatory neurons and *N*_*I*_ inhibitory neurons, receiving inputs from *N*_*X*_ external excitatory neurons. The dynamics of the firing rate of the *i*-th excitatory (*a* = *E*) or inhibitory (*a* = *I*) neuron, 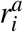, is described by

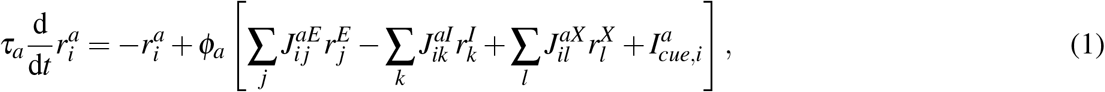

where *ϕ*_*a*_ is a sigmoidal activation function, and 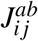 are the synaptic weights from neuron *j* of population *b* = *E, I, X* to neuron *i* of population 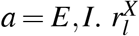 are the firing rates of external neurons.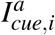 is an external input that provides a cue at the beginning of sequence retrieval to neuron *i* (see below). *J*^*ab*^ is sparse, and is either random or stores *S* random sequences via a Hebbian rule. For simplicity, this study considers sequences that are discrete in time, though continuous sequences could also be stored [15]. The connectivity matrix after storing *S* sequences is given by

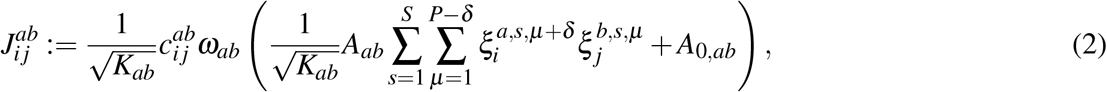

where *K*_*ab*_ is the average number of synaptic inputs from population *b* to a neuron in population 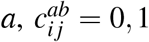 are i.i.d. Bernoulli random variables, *A*_*ab*_ is the Hebbian learning strength, *A*_0,*ab*_ is the baseline weight, and *ω*_*ab*_ is a threshold-linear function enforcing the sign of the weights from *b* to *a*. Using softplus, *ω*_*ab*_(*x*) = *m*^−1^ ln(1 + exp(*mx*)), with large *m* values (*m* = 30, 50) leads to qualitatively similar results. *δ* = 0 for a temporally symmetric Hebbian (TSH) rule and *δ* = 1 for an asymmetric (TAH) rule. For any sequence *s*, TAH plasticity associates the *µ*-th pattern, *ξ*^*s*,*µ*^, with the next pattern, *ξ*^*s*,*µ*+1^. TSH plasticity associates *ξ*^*s*,*µ*^ with itself. For simplicity, we assume *ξ*^*s*,*µ*^ ∼ *N*(0, 1) are independent for all *s* and *µ*.

When *A*_*ab*_ = 0 for the connectivity from population *b* to *a*, there is no learning or plasticity, and the corresponding connectivity is sparse and random, i.e.

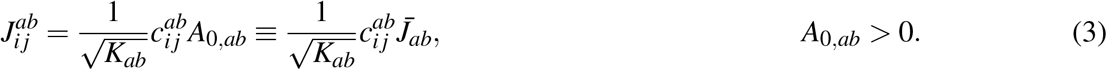

Notice that *J*^*II*^ (see below), *J*^*EX*^ and *J*^*IX*^ are random throughout this study.

### 2.2 Sequence Retrieval

A recent study [15] demonstrated that an E-I network with a specific structure can store and retrieve sequences using the TAH rule described by Eq. 2 in E→E connections. We first verified that sequences can be stored and retrieved using TAH rule in E→E connections (*A*_*EE*_ *>* 0 for *J*^*EE*^, Eq. 2) in a balanced E-I network described by Eq. 1 (Fig. 1A). This model resembles the E-I network in [15], but ours has I→I connections and dynamically reaches E-I balance. For convenience, we call the model with only E→E plasticity “EE model”. Upon storage of the sequences, the network is provided with an external input, 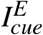, proportional to the first pattern of the retrieved sequence for 20 ms (see Section 4.3). The excitatory and inhibitory populations both displayed sequential peaks of neuronal firing rates, while the population means and standard deviations of the firing rates and synaptic inputs (defined as the sum of all inputs to a neuron; Eq. 18) remained stable (Fig. S1A-D).

**Figure 1:**
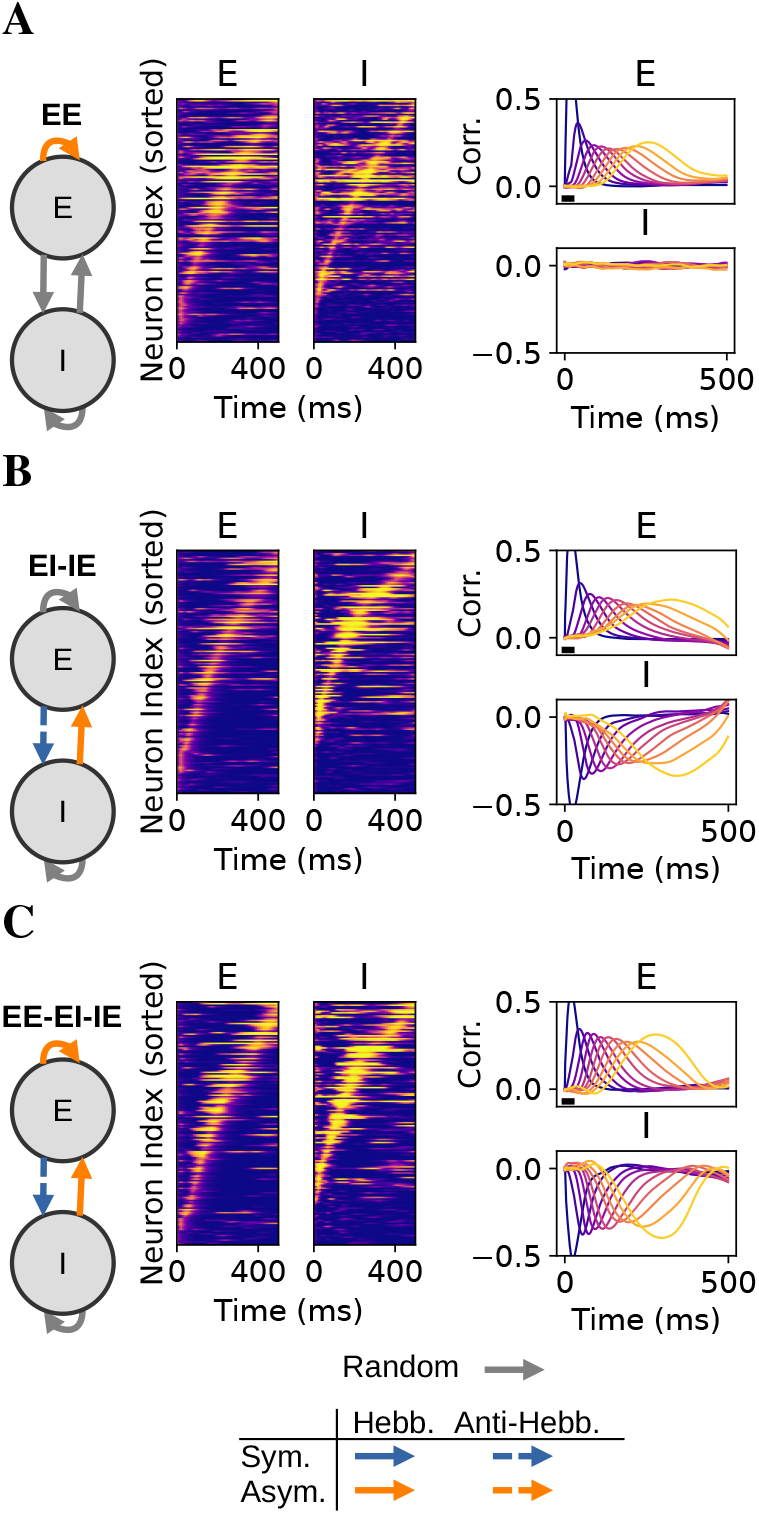
Sequences can be stored and retrieved in the absence of E→E plasticity. (A) Left, schematic of the EE model. Middle, activity of excitatory (left) and inhibitory (right) populations vs time during retrieval of one of the stored sequences. For clarity, only a random subset of *n* = 2000 active neurons (temporally averaged neuronal firing rate *>* 1 Hz) is shown. Brighter (darker) colors indicating higher (lower) firing rates. Right, correlation between the firing rates of excitatory (top) or inhibitory (bottom) neurons and the patterns in the retrieved sequence. Brighter colors indicate later patterns. The initial pattern of the sequence is presented as an external input from 1 ms to 20 ms (thick black bar). (B) and (C) as (A) but for EI-IE and EE-EI-IE model, respectively. Parameters in Table 3.

To characterize retrieval dynamics, we calculated the overlaps 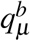 between the neuronal firing rates and patterns in the retrieved sequence

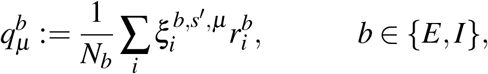

and the Pearson correlations between both (overlaps divided by the standard deviations of the firing rates and patterns). Both quantities describe the linear relationship between sequence patterns and neural activity. Using dynamical mean-field theory (DMFT), it is possible to derive the overlap dynamics (see Section 4.1):

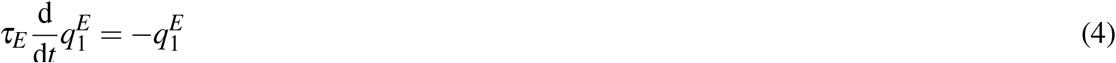

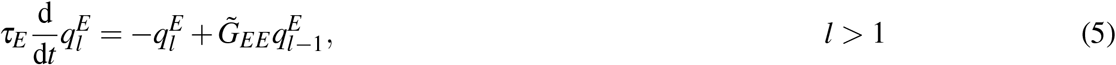

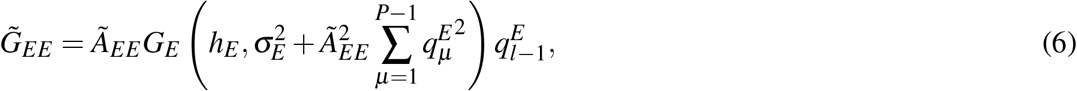

where *Ã*_*EE*_ is a parameter describing the strength of learning, and *G*_*E*_ is a function of the mean and standard deviation of the total synaptic inputs to excitatory neurons, *h*_*E*_ and *σ*_*E*_, respectively (see Section 4.1). In our analysis, we assume the overlap with the first pattern, 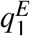, is set by an initial condition correlated with the first pattern and decays exponentially (Eq. 4). Eq. 5 shows that the overlap with a pattern *l >* 1 is driven by the overlap with its previous pattern, resembling a “delay line” system. We confirmed our DMFT approximation matches qualitatively the simulations of network models (Fig. S6F).

As predicted by Eq. 5, during retrieval in the EE model, the correlations peaked successively over time, in the same order as the corresponding patterns (Fig. 1A). This indicate that the sequence is retrieved in the correct order. Further, peaks of neuronal firing rates are narrower at the beginning of the sequential activity than the later part (Fig. S1E), and correlation peaks are wider for later patterns as well (Fig. S1F), consistent with [15] and experiments [1, 44, 47].

### 2.3 Sequences can be Stored and Retrieved in the Absence of Recurrent Excitatory Plasticity

We then asked weather sequences can be stored in synaptic connections that involve inhibitory neurons, because emerging evidence suggests that diverse forms of plasticity can exist in E→I [4, 53] and I→E connections [9, 23, 26] across different brain areas [14, 32, 49, 57]. I→I plasticity may also exist, but the direct experimental support is scarce [32, 57], and therefore not considered in this study.

We found that networks can store and retrieve sequences with both E→I and I→E plasticity, as long as the E→I→E pathway effectively implements TAH learning (Fig. 1B and Fig. S5). This means (1) either E→I or I→E is Hebbian and the other is anti-Hebbian, and (2) one of E→I or I→E is temporally asymmetric, while the other is temporally symmetric. For example, a possible scenario is that E→I connections are shaped by a temporally symmetric anti-Hebbian (TSAH) rule (*A*_*EI*_ *<* 0 for *J*^*EI*^, Eq. 2), and I→E connections are shaped by a TAH rule (*A*_*IE*_ *>* 0 for *J*^*IE*^). We referred to this set of models as “EI-IE” models.

EI-IE models displayed similar sequential activity and retrieval dynamics as the EE model when the initial cue 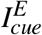 was set to be proportional to the first pattern of a stored sequence *s* (Fig. 1B and Fig. S2). We set 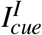 to zero, because 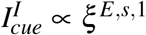 is sufficient to drive sequence retrieval. In both simulations and theoretical analysis, we assumed inhibition is much faster than excitation (*τ*_*I*_ ≪ *τ*_*E*_). It is possible to derive the overlap dynamics for the EI-IE model (Section 4.1),

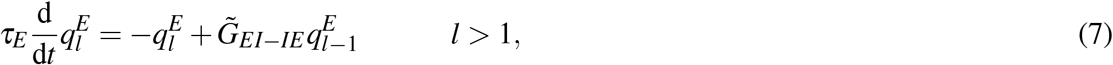

where 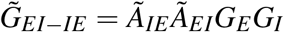. Compared to 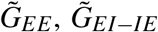 depends in addition on the mean and standard deviation of total synaptic inputs to inhibitory neurons, *h*_*I*_ and *σ*_*I*_, but this does not qualitatively affect the overlap dynamics, in which 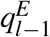drives 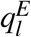 .

We also investigated “EE-EI-IE” models, in which both TAH pathways exist, i.e. E→E and E→I→E. The dynamics does not qualitatively differ from that of the EE and EI-IE models, as shown by simulations (Fig. 1C and Fig. S3) and the overlap dynamics assuming fast inhibition (Eq. 8).

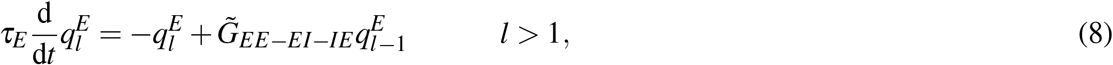

where 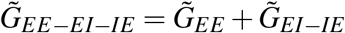.

In short, similarly to the previously studied EE model, both EI-IE and EE-EI-IE models can store and retrieve sequences. The three models have qualitatively similar and sequential activity, and all function like a “delay line” system during retrieval. Therefore, recurrent excitatory plasticity is not necessary for sequence storage and retrieval, as long as the E→I→E pathway is effectively TAH. Networks storing much longer sequences via different combinations of plasticity rules also have similar dynamics of sequence retrieval (Fig. S4), suggesting this is a general observation.

### 2.4 EI-IE Model can achieve higher storage capacities than both EE and EE-EI-IE Models

We have shown in the previous section that sequences can be stored in different ways in recurrent E-I networks, using either exclusively E-E plasticity, an appropriate combination of E-I and I-E plasticity, and a combination of EE, EI and IE plasticity. We next ask which of these scenarios maximize storage capacity.

We define the memory load *α*^(*m*)^ of a model *m* ∈ {EE, EI-IE, EE-EI-IE} as

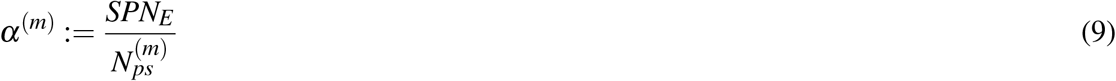

where the numerator is proportional to the total amount of information stored in the excitatory network, while 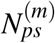 is the total number of plastic synapses. This definition is a generalization of measures of information storage used in previous studies [3, 12, 16, 21] to a E-I networks with various plasticity scenarios. It measures how much information is stored in the network per plastic synapse, and is expected to remain of order 1 in the large *N* limit.

To find the storage capacity of a model, 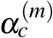, i.e. the maximal memory load at which sequences can still be retrieved successfully, we observe that when 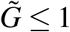, the maximal overlap with a pattern 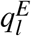 is smaller than the maximal overlap with the previous pattern 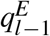. In this case, long sequences cannot be fully retrieved, since overlaps progressively decay to zero during retrieval. For sufficiently large *α*^(*m*)^, 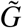 is a decreasing function of *α*^(*m*)^. Thus, either 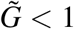 for all *α*^(*m*)^ and the capacity is zero, or the capacity 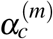 is given by the larges value of *α*^(*m*)^ for which 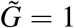[15] (Section 4.2). In both simulations of the networks and solutions to the dynamical mean field equations (DMFEs) with corresponding parameters, the maximal correlations with the later patterns of the retrieved sequence sharply drop to zero when the memory load of a model is larger than its storage capacity found using this method, but not when it is smaller (Fig. S6A-E). This indicates that our method based on the values of 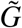 gives a good estimate of the actual storage capacity of a network. Interestingly, for certain parameters of the EI-IE model, solutions to the DMFEs predict much higher peak correlations compared to the simulations. Increasing the network size cannot reconcile the difference, suggesting that it is not simply due to the finite size effect. A possible reason is chaotic dynamics (Fig. S7).

We performed grid searches over the learning strengths *A*s, baselines of synaptic weights *A*_0_s^1^, external input weights 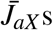, and the inhibitory weight ratio 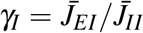. We chose the ratio *γ*_*I*_ but not 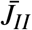 as a parameter because EI balance is strongly affected by the ratio between the two, but not their magnitudes (unless both are too small or too large). Also, in the EI-IE and EE-EI-IE models, 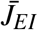 grows with *α*(*m*). Searching over *γ*_*I*_ allows 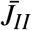to grow with 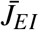 and is more efficient than searching over a wide range of *J*_*II*_. Our grid search showed that the EI-IE model has higher optimal capacity than the other two models (Fig. 2A; 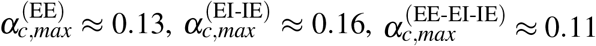, where 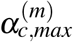 is defined as max{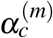}). Furthermore, the grid search suggests that the distribution of storage capacity of EI-IE model seems to be more spread out across all searched parameter sets, compared to the other two models (Fig. 2B). To test the significance, we define relative capacity of a parameter set as the ratio between its storage capacity and the optimal capacity of the model, i.e. 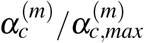. We found that the EI-IE model is more likely to have a greater relative capacity than EE-EI-IE model than EE model (*p <* 10^−5^, one-sided Brunner Munzel test). In other words, EI-IE model may have a larger range of parameters that allow higher capacity than the other two models.

**Figure 2:**
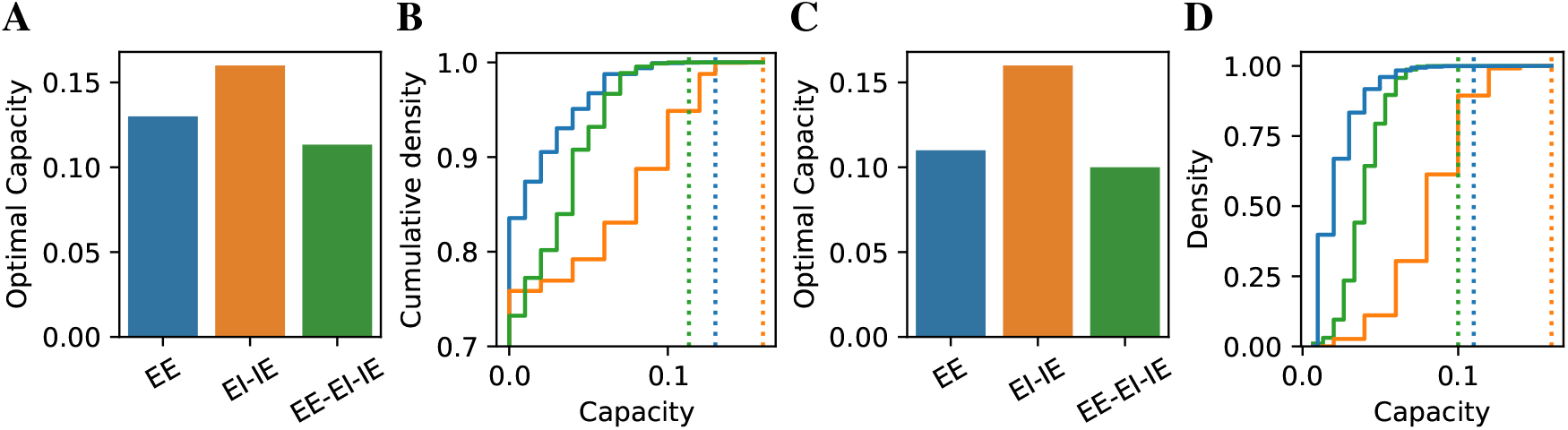
Grid search over the three models shows that EI-IE model can achieve a higher storage capacity than EE and EE-EI-IE models. (A) The optimal capacity for each model. (B) The cumulative distribution of capacities for each model, across all sampled parameter sets. (C-D) as (A-B), but parameter sets leading to unrealistic firing rates are excluded. We consider [1-10] Hz as realistic average excitatory firing rates, and [10-30] Hz as realistic average inhibitory firing rates.

It is important to ensure that the optimal capacity solutions are consistent with experimental observations, so we next restricted our analysis to parameter sets that lead to realistic stationary mean firing rates (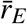 between 1 and 10 Hz, and 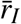 between 10 and 30 Hz). The optimal capacity of the EI-IE model among such sets remained unchanged, though the optimal capacities of EE and EE-EI-IE model slightly decreased (Fig. 2C; 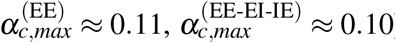). The EI-IE model with realistic stationary mean firing rates is again more likely to achieve higher capacity relative to the optimum than the other two models (Fig. 2D; *p <* 10^−5^ one-sided Brunner Munzel test).

To gain more insights on how high- and low-capacity solutions differ within and between models, we investigated the distributions of grid search parameters, mean synaptic weights, and stationary statistics of population activities for different solutions at their storage capacities. We defined high-capacity solutions as those with memory load 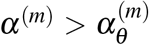, and low-capacity solutions as those with 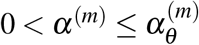, where 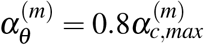 is a model-specific threshold. High-capacity solutions in general have large learning strengths (Fig. S8), as expected intuitively. We also observed a strong preference of negative negative *A*_0_ for plastic connections, except in the EE-EI-IE model.

Importantly, we observed that the excitatory weight ratio 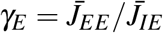is almost strictly smaller than the inhibitory weight ratio 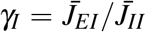for all non-zero capacity solutions (Fig. 3A). Yet, the mean E→I synaptic weights is almost strictly larger than I→I synaptic weights only in high-capacity solutions (Fig. 3B). This suggests that high-capacity solutions are strongly aligned with the E-I balance conditions [51, 52]. As a result, the mean inhibitory firing rate is larger than the mean excitatory firing rate (Fig. 3C). Interestingly, we found that the mean synaptic input to excitatory neurons is more negative (inhibitory) than the mean synaptic input to inhibitory neurons (Fig. 3D). This might be an effect of EI balance, since inhibition has to be stronger than excitation in balanced networks. The standard deviation of the input to excitatory neurons is also larger in magnitude (Fig. 3E), ensuring that at least a fraction of the excitatory neurons are active. However, the result is a larger *G*_*I*_ than *G*_*E*_ (Fig. 3F), because in active states, *G* decreases when the magnitudes of mean and standard deviation of the synaptic inputs increase (Eq. 29).

**Figure 3:**
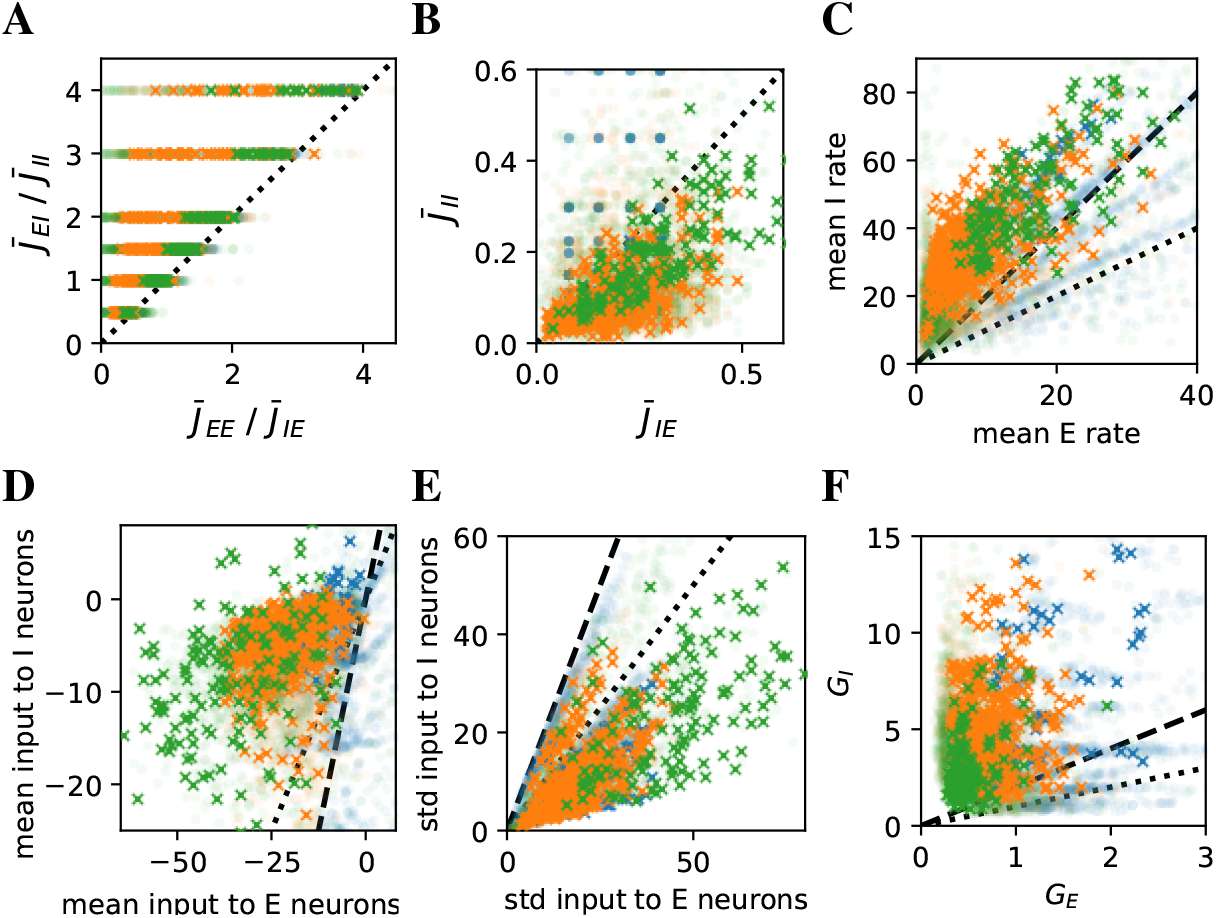
High-capacity solutions strictly obey the E-I balance conditions and have more negative and variable synaptic inputs to excitatory neurons than to inhibitory neurons. Different measures for excitation (horizontal axis) and inhibition (vertical axis) are plotted against each other. Blue, EE; orange, EI-IE; green, EE-EI-IE. Each symbol corresponds to a solution (a parameter set). Crosses indicate solutions with capacities larger than 80% of the model’s optimal capacity, and transparent circles indicate solutions with small, but non-zero capacities. Zero-capacity solutions are excluded from the plots. Dotted line, identity *y* = *x*; dashed line, *y* = 2*x*. (A) Ratios between mean excitatory synaptic weights plotted against inhibitory synaptic weights. (B) Mean E→I weights plotted against mean I→I weights. (C) Mean firing rates. (D-E) The means (D) and standard deviations (E) of the synaptic inputs to excitatory or inhibitory neurons. (F) Overlap gain functions of inhibitory or excitatory populations.

All together, the grid search over parameters shows that the EI-IE model has a higher optimal capacity than EE and EE-EI-IE models, even when physiological constraints on mean firing rates are considered. High-capacity solutions closely follow the E-I balance conditions and have more negative and variable synaptic inputs to excitatory neurons than inhibitory neurons.

### 2.5 Inhibitory Plasticity Improves Robustness to Initial Cue Perturbation and External Noise

We have investigated so far how and why models with and without inhibitory plasticity may have different storage capacity, in the absence of any noise during retrieval. In realistic conditions, whether retrieval is successful is expected to depend on external noise and the degree of correlation of the initial cue with the first pattern in the sequence. It is thus crucial to explore how these factors affect retrieval, and how robust are the different models to external noise and initial cue perturbation.

To study whether inhibition enhances robustness of sequence retrieval, we varied initial cue perturbations and external noise strengths, and compared the three models. We initialized the excitatory neurons with a perturbed version of the first pattern of the sequence to be retrieved, 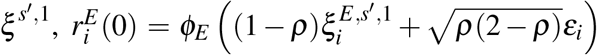, where 0 ≤ *ρ* ≤ 1 is the perturbation strength, and *ε*_*i*_∼*N*(0, 1) are i.i.d. random variables. For the external noise, we assumed 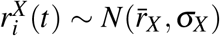 with fixed 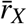 and varied the coefficient of variation (CV), defined as 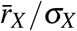, over simulations.

We ran multiple simulations (*n* = 20) of the three models with similar sets of parameters (Table 7 and 8) that produce physiological stationary firing rates (5 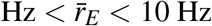 and 15 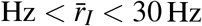), and memory load close to their capacity (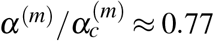 for Table 7 and 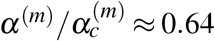 for Table 8. Notice that the ratios cannot be exactly the same for different models, because the sequences are composed of a discrete number of patterns). At theindividual neuron level, peaks and valleys in the firing rates become less distinguishable when the initial cue perturbation *ρ* (Fig. 4A) or external noise CV (Fig. 4B) increases. Strong initial cue perturbation and external noise also prolong or diminish peaks that originally exist without initial cue perturbation and noise, or create new peaks in the firing rates. At the population level, the maximal correlations with sequence patterns decrease when the initial cue perturbation and external noise CV increase (Fig. 4C-D).

**Figure 4:**
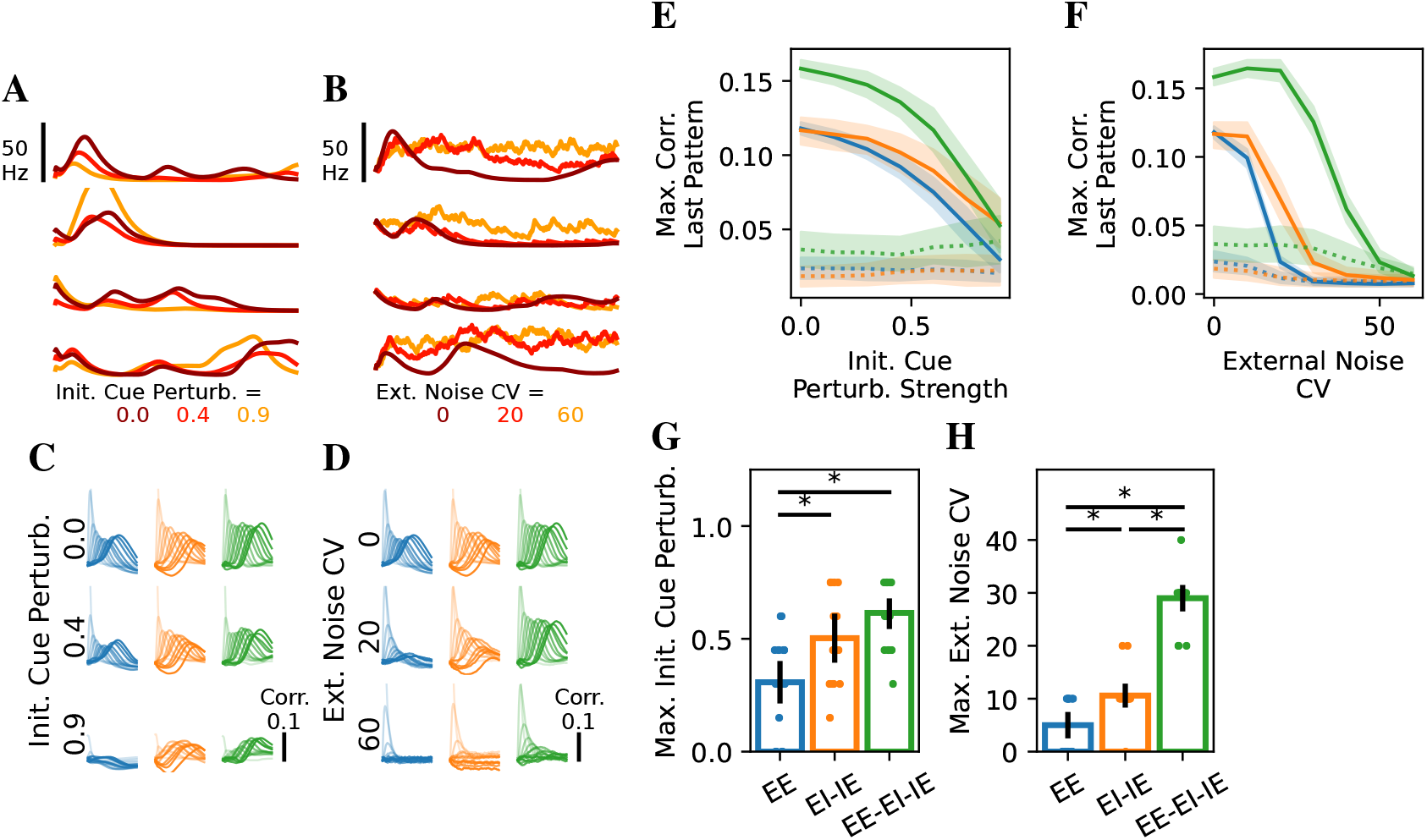
Inhibitory plasticity enhances robustness to initial cue perturbation and external noise. (A) Firing rates of four example excitatory neurons of the EE model for different strengths of initial cue perturbation. Brighter color indicates stronger perturbation (weaker cue). (B) As (A), but for different external noise CV. For (A-B), the disruptions in neuronal firing rates of the EI-IE and EE-EI-IE models are not visibly different compared to those of the EE model, so only neurons from the EE model are plotted. (C-D) Correlations with the patterns in the three models (EE, blue; EI-IE, orange; EE-EI-IE, green) for different levels of initial cue perturbation (C) or external noise CV (D). For (A) and (C), the external input is constant, and for (B) and (D) the initial cue is not perturbed. (E-F) Maximal correlation with the last pattern of the retrieved sequence as functions of initial cue perturbation (E) or external noise (F). Solid/dotted lines: Maximal correlation with the last pattern the retrieved/other sequences, respectively. The lower and upper boundaries of the shaded areas around the curves mark 1 s.t.d. around the mean. (G-H) The maximal initial cue perturbation *ρ* (G), or and the maximal external noise CV (I) each model could tolerate until retrievals failed. Error bands and bars denote the 95% confidence interval, and asterisks indicate statistical significance. Parameters are described in Table 7.

We developed multiple measurements to quantify the retrievals under noise and perturbation (Section 4.5). A successful retrieval can require many factors, such as significant correlations with the retrieved sequence, and distinct and correctly ordered peaks of correlations. We chose the following criteria for successful retrieval:

1. The maximal correlation with the last pattern should be larger than 0.1;
2. The peaks of correlations are in the correct order; and
3. The correlation peak definiteness (averaged ratio between a peak at a certain time *t* and the maximum among the correlations at *t*) is larger than 0.95.

The maximal correlation with the last pattern decays gradually when the initial cue perturbation or external noise becomes stronger (Fig. 4E-F and Fig. S10E-F). The other two criteria, the percentage of retrieval with correct peak order and correlation peak definiteness, remain relatively constant until the perturbation and noise are large (Fig. S11). Other measurements we devised also agree with the three criteria (Sec. 4.5). All of these criteria are failed when they are applied to unretrieved sequences for all perturbation and noise strengths.

By applying these criteria, we found that EI-IE and EE-EI-IE models are more robust to initial cue perturbation and external noise than the EE model (Fig. 4G-H and Fig. S10G-H). EI-IE and EE-EI-IE models can have significantly larger averaged maximal initial cue perturbation strength (*p <* 0.005, two-sample t-tests). EE model can tolerate smaller external noise CV on average than the other two models (*p <* 0.01, two-sample t-tests). We observed no consistent differences in maximal perturbation strength between EI-IE and EE-EI-IE models. However, sequences can be retrieved in EE-EI-IE model with stronger external noise than EI-IE model (*p <* 10^−5^, two-sample t-test).

## 3 Discussion

In this study, we analyzed the impacts of inhibition and inhibitory plasticity to sequence storage and retrieval in a sparsely connected E-I network with different combinations of Hebbian plasticity rules. Using both simulations and a mean field approach, we found that networks with diverse forms of plasticity rules in both E→I and I→E connections (EI-IE model) can store and retrieve sequences, as long as the E→I→E pathway is effectively temporally asymmetric and Hebbian. The characteristics of the sequential activity in the EI-IE model do not differ qualitatively from a network with TAH plasticity in only E→E connections (EE model), as well as a network with E→E, E→I, and I→E plasticity (EE-EI-IE model). Grid searches over learning-related parameters showed that EI-IE model has the largest optimal capacity, and are most likely to reach a high capacity relative to the optimum. Moreover, inhibitory plasticity also improved robustness to external noise and initial cue perturbation during retrieval of sequences. In conclusion, this study suggests new computational roles for inhibitory plasticity in enhancing sequence storage capacity and robustness of sequence retrieval.

How sequences are learned and retrieved has been studied by theorists for decades. However, few studies have addressed the roles of inhibitory plasticity apart from maintaining E-I balance, in spite of the fact that multiple forms of inhibitory plasticity have been found experimentally [4, 9, 14, 22, 23, 26, 32, 49, 53, 57], and suggested to be critical in sequence learning [1, 4, 22, 29, 49]. Why do various forms of inhibitory plasticity exist, and how do they affect sequence learning? We take a step towards answering these questions by studying multiple scenarios involving inhibitory plasticity rules. We showed that models with no E→E plasticity can produce sequential activity, provided plasticity in both E→I and I→E pathways exist. In contrast with widely studied synfire-chain models [5, 11, 13] and binary networks with Hebbian plasticity [15, 16, 24, 25, 38], the spatiotemporal dynamics of our models can match a diverse sets of experimental observations [1, 44, 47]. While models storing sequences in E→E connections only [5, 11, 13, 15] and in I → I connections only[36] have been implemented, it remained unclear how models with plasticity in excitatory and/or inhibitory connections differ. Analyzing models with various forms of plasticity rules, we discovered multiple benefits of inhibitory plasticity in enhancing storage capacity and robustness or retrieval.

Our findings are closely related to a few theoretical works that explored the impact of inhibitory plasticity on learning non-temporal patterns. Vogels *et al*. [54] demonstrated that symmetric Hebbian inhibitory plasticity can lead to global E-I balance and sparse asynchronous irregular firing in recurrent neural networks. This regime has been suggested to be critical for high capacity and noise robustness in associative memory networks [43]. Mongillo *et al*. [34] further showed that networks with effectively Hebbian plasticity in connections involving inhibition can store many more binary patterns than a model that relies purely on Hebbian plasticity in EE connections, and suggested that plasticity in inhibitory, rather than excitatory, connections might maintain information over long periods in networks with volatile connectivity[35]. Another recent study [26] showed that asymmetric plasticity from SOM-INs to pyramidal cells stabilize assembly formation, helping assemblies to be more robust to perturbation. Thus, inhibitory plasticity has been suggested to benefit memory storage capacity and retrieval robustness in models that store independent patterns. We have generalized this idea to sequences, and used a novel analysis to explain why storing sequences in E→I→E pathway can have a higher optimal capacity.

Interestingly, we also observed from grid searches that models with capacity close to their optima tend to have negative weight offsets *A*_0_ in plastic synapses (Fig. S8) In these cases, many weights are zero due to the sign constraint, unless the pre- and postsynaptic patterns are strongly correlated. This is consistent with previous findings in networks with sign-constrained synapses [6, 7, 8, 10, 43], where a large fraction of zero synaptic weights (referred to as ‘silent’ or ‘potential’ synapses) was found to be necessary for optimizing storage capacity robustly. This results in an effectively greater sparseness than the specified connectivity probability. An interesting future direction would be to systematically quantify how sparseness affects the storage and retrieval of sequences.

Compared to previous models [15, 16], our optimal storage capacities are lower. In sparsely connected networks without sign constraint that stores sequences using bilinear TAH rule, the optimal storage capacity is ∼0.2-0.3 for binary neurons [16] and ∼0.4-0.5 for analog neurons [15]. In contrast, the optimal storage capacities of the models we analyzed is ∼0.1-0.2. A few reasons could cause this discrepancy. First, it is well known that sign constraints on synaptic weights lead to lower storage capacity [2]. Second, maintaning E-I balance provides additional constraints that might further decrease the optimal capacity. Our simulations indicate that in some parameter regimes, our networks exhibit chaotic dynamics (Fig. S7), which could lead to lower correlations with the retrieved patterns and a smaller capacity. A detailed theoretical characterization of the chaotic dynamics is beyond the goal of this study. Future work should investigate whether these or other factors are responsible for the reduction in storage capacity compared to unconstrained networks.

Our results depend on how memory load and capacity are different, or in other words which “cost function” is optimized. We have defined the capacity as the maximal number of patterns that can be stored per plastic synapse, ignoring non-plastic synapses in the network. This corresponds to maximizing a cost function with an energetic term in which plastic synapses cost much more energy than non-plastic synapses [33, 40, 41]. However, if energy and resources are less important than the absolute amount of information stored, then one should replace the denominator of Eq. 9 by the total number of synapses. Then, the EE-EI-IE model has the highest optimal capacity, since it has the most plastic synapses. In both cases, adding plasticity to both EI and IE connections is beneficial to capacity.

The networks we investigated include a single homogeneous inhibitory population. Yet, various types of inhibitory neurons exist in different brain structures and display diverse cellular properties and connection profiles [14, 32, 49, 57]. It will be therefore important to explore the impacts of heterogeneity of inhibitory neurons in future studies.

## 4 Methods

### 4.1 Dynamical Mean Field Equations under Sequence Retrieval

The network model described by Eq. 1 is high-dimensional. We can apply the dynamical meanfield theory (DMFT) to develop a low-dimensional description of the behavior of the network [15, 45, 48]. Using DMFT, the dynamics of the network can be reduced to a small number of variables characterizing the statistics of network activity, such as the means, variances, and overlaps between the neural activity and the stored patterns.

DMFT is usually performed in networks where weights are a linear sum of patterns [15, 48]. To deal with non-linearity, we have to use a linear approximation of the weights. By assuming that during the retrieval the neural activity is only non-trivially correlated with the retrieved sequence, it is convenient if the retrieved sequence can be separated from the others. Under the large *K* limit, we can expand the plastic weight matrices for any retrieved sequence *s*^′^ if the contribution of a single sequence is small compared to the rest of the argument of *ω*. Possible scenarios include

1. The load *S*(*P* − 1) is small, and the offset *A*_0,*ab*_ is positive.
2. Many sequences are learned, i.e. *S* ≫ 1.

This gives

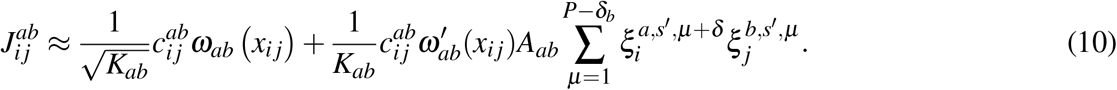

where

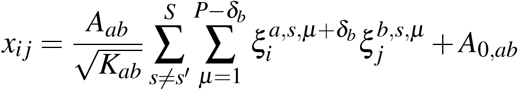

For large *S* × *P, x*_*ij*_ can be approximated as a Gaussian noise with mean *A*_0,*ab*_ and variance 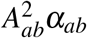, where

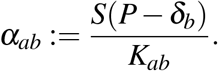

Notice that *α*_*ab*_ is the “load” of the connections from population *b* to population *a*, and is *not equivalent to* the memory load *α*^(*m*)^ of the network, except for the EE model *α*_*EE*_≡*α*^(EE)^. With this approximation,

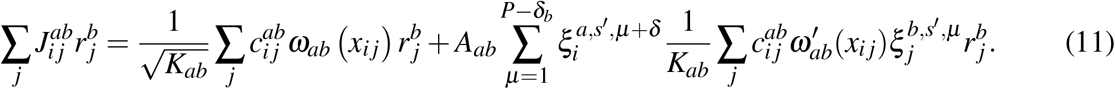

Under the large *N* limit, the input from population *b* to *a* can be approximated by a Gaussian random variable. That is,

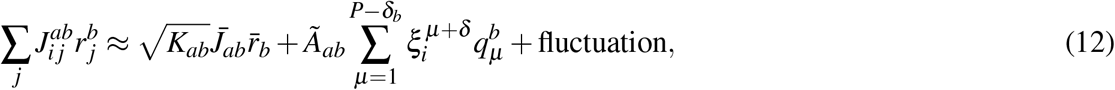

where 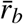 is the mean rate of population *b*, and we define the overlap between neural activity and pattern *µ* of population *b* as

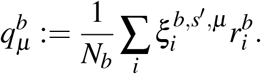

For convenience, we drop *s*^′^ for the retrieved sequence in the notation of overlap *q*’s. 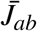and *Ã*_*ab*_ are the scaled mean synaptic weight (“scaled” because the actual mean synaptic weight is 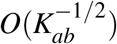) and the effective learning strength, respectively.

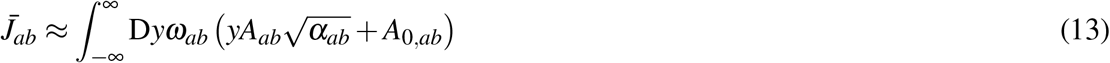

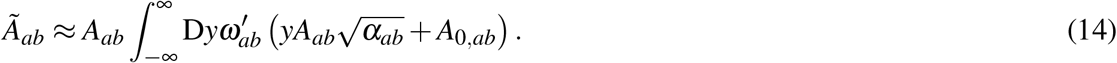

Notice that here *ω*^′^ is the derivative of *ω*.

For large *S*(*P* − 1), if *A*_0,*ab*_ ≈ 0 and *ω*_*ab*_ is nearly a threshold-linear function,

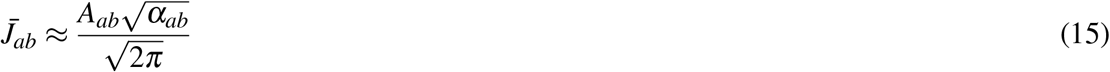

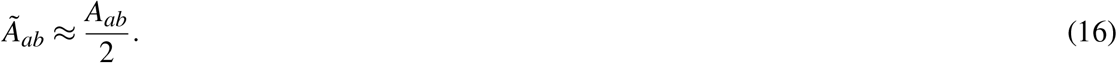

If *J*^*ab*^ is not plastic, the mean is similar to Eq. 12, except that the second term of Eq. 12 is zero due to *Ã* = 0.

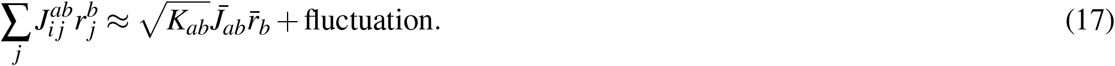

Finally, the mean of the synaptic input to population *a* is

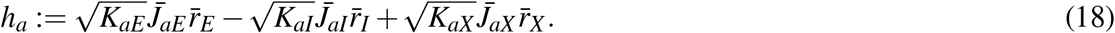

In the large *K* limit, the mean excitatory and inhibitory inputs must balance, such that *h*_*a*_ ∼ *O*(1) [51, 52]. We have the following balanced equations for *a* ∈ {*E, I*}:

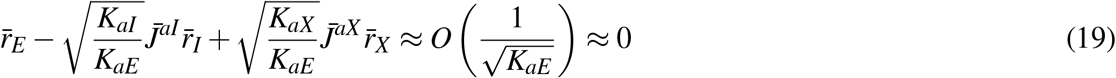

For simplicity, we can set *K*_*aE*_ = *K*_*aX*_, and the ratio *K*_*aI*_*/K*_*aE*_ is (approximately) cancelled by the difference between 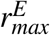 and 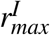. The conditions for E-I balance could be derived [51, 52]:

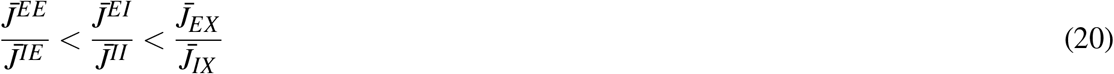

Further, enforcing 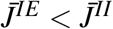 will eliminate solutions that saturate at the maximal firing rates [51, 52]. However, in the cases where 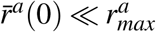, the stationary states reached are likely to be nearly silent states. Further, in our models with inhibitory plasticity, this condition may not always be necessary due to the effect of inhibitory overlaps. Therefore, we do not strictly require 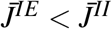 in our parameters.

Next, we need to derive the fluctuation term. We assume that the external input is not correlated with the feedback or recurrent inputs, and the activities of two different neurons are nearly uncorrelated because each neuron receives from many other neurons.

Under the large *K* limit, the parts of order 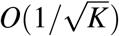 and higher are dropped in the calculation of autocovariance (that is, the covariance between the first and second terms of Eq.10). Consequently, the fluctuation comes from the sparse connections whose synaptic strengths are shaped by sequences that are independent of the current one. The variance of the total input to population *a* is

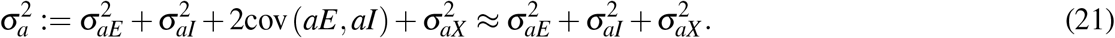

For the autocovariance of an input, *ρ*_*ab*_(*s, t*), in the plastic case,

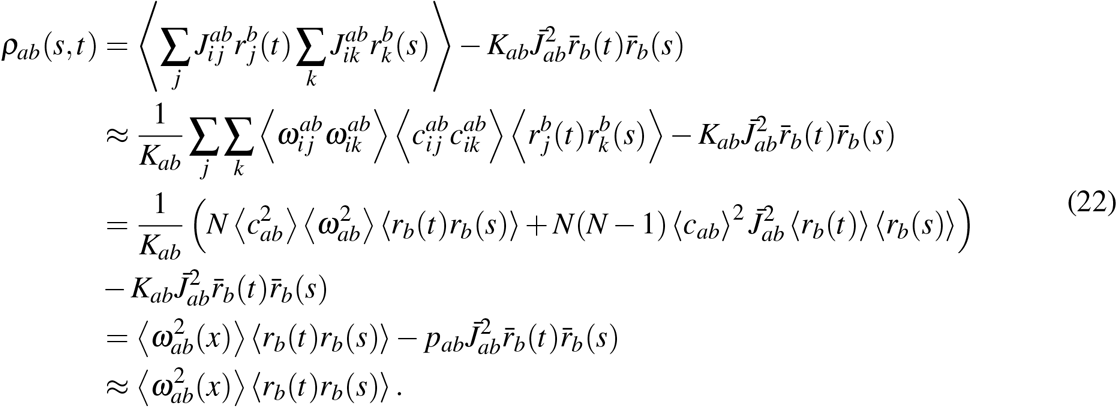

And we define the variance as

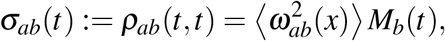

where 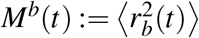. Notice that if *A*_0,*ab*_ ≈ 0 and *ω*^*ab*^ is (or close to) a threshold-linear function,

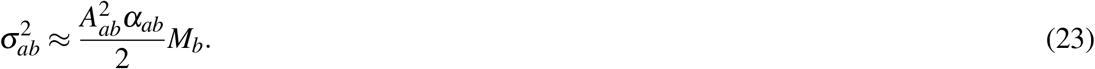

If the input connectivity is not plastic,

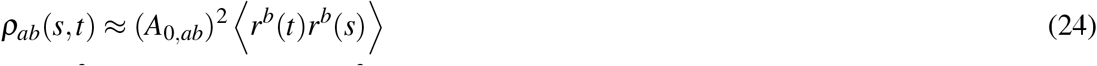

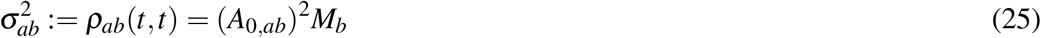

Finally, we need to derive the overlap dynamics. As an example, consider the model with only TAH plasticity in E→E connection (EE model):

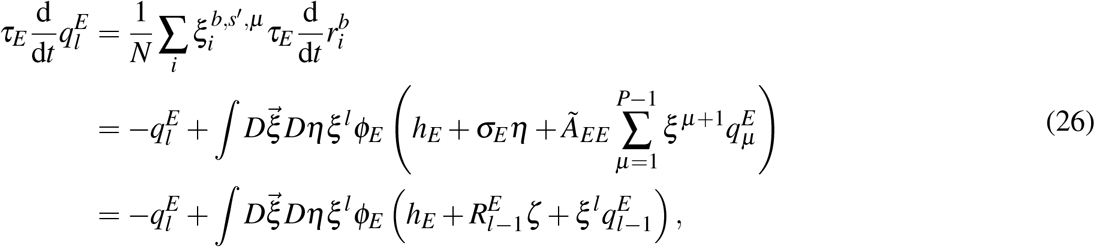

where 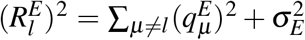. In this case, the overlap with the first pattern in the sequence (*l* = 1) always decays exponentially, i.e.

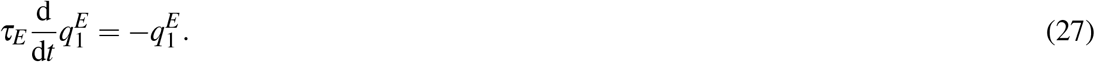

For *s >* 1, apply the change of variables [15]

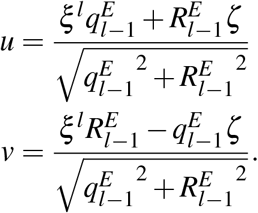

Eq. 26 becomes

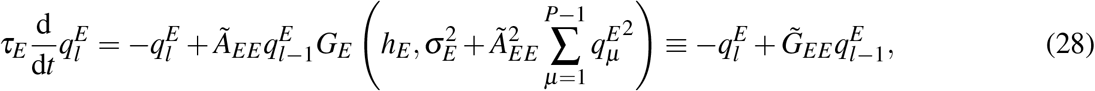

where *G*_*a*_(*c, σ* ^2^) := *σ*^−2^*∫Dvvφ*_*a*_(*σv* + *c*). For sigmoidal activation function,

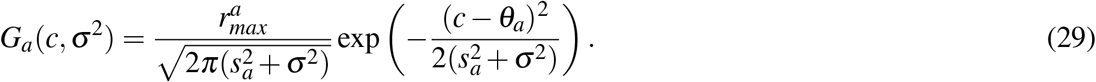

Following the example of EE model, we can also derive the overlap dynamics for a model with E→I and I→E plasticity (EI-IE) model. Here, we chose E→I to be TSH and I→E to be TAH, but the reverse is similar.

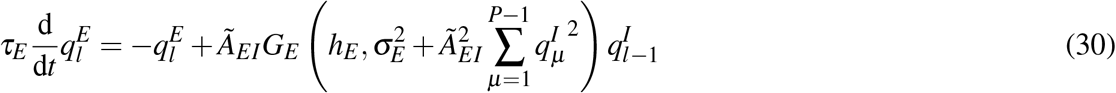

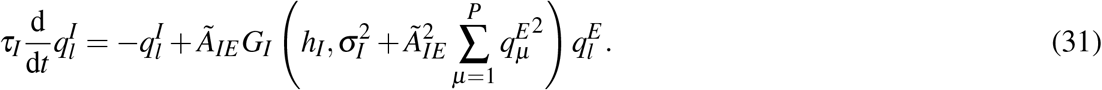

By assuming *τ*_*I*_ ≪ *τ*_*E*_, *q*^*I*^ ≈ _*IE*_*G*_*I*_*q*^*E*^, and

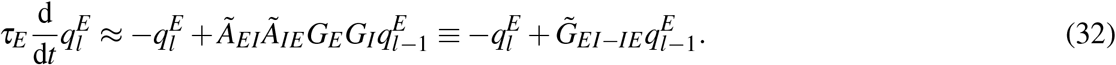

The overlap dynamics for a model with TAH E→E, TSH E→I and TAH I→E plasticity (EE-EI-IE) model, after assuming *τ*_*I*_ ≪ *τ*_*E*_, is

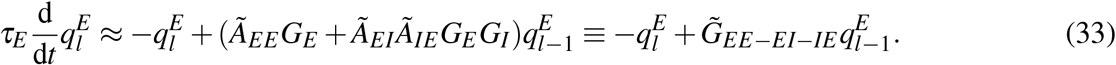

Thus, for models with an effectively temporally asymmetric feedback loop, the overlap with a pattern is driven by the overlap with its previous pattern.

Finally, we will give the general dynamical mean field equations (DMFEs). For a one-population network storing discretized sequences with bilinear rule, the equations were derived previously [15]. Here, we generalize them to a balanced E-I network. As mentioned above, the dynamics of the mean rates and firing rate variances can be approximated using the first and second order statistics of the inputs (Eq. 18 and Eq. 21). We can derive the s f-consistent equations of population mean 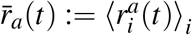, second moment 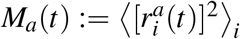 and (uncentered) covariance 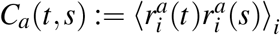.

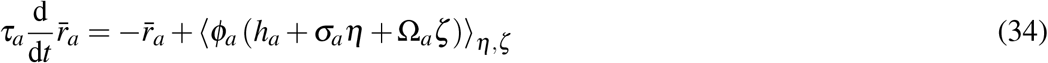

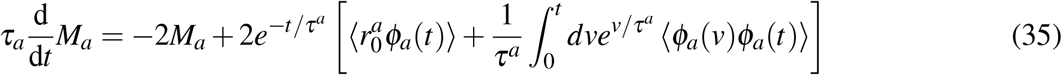

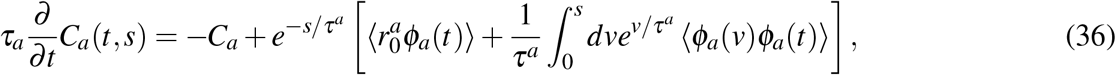

where *h*_*a*_ and *σ*_*a*_ are given by Eq. 18 and Eq. 21, respectively, and

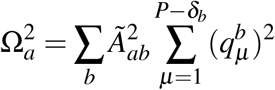

is the variance of the fluctuation due to sequence retrieval.

The derivations of *M*_*a*_ and *C*_*a*_ use the solution to individual firing rate:

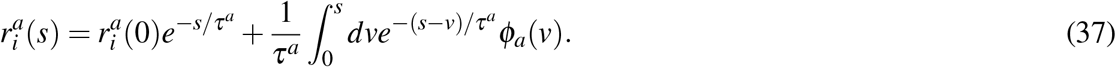

For convenience, we use abbreviations *φ*_*a*_(*t*) ≡ *φ*_*a*_ (*h*_*a*_(*t*) + *σ*_*a*_(*t*)*η*_*t*_ + Ω_*a*_(*t*)*ζ*_*t*_) and similarly for *φ*_*a*_(*v*). The ensemble average ⟨*φ*_*b*_(*v*)*φ*_*a*_(*t*)⟩ is

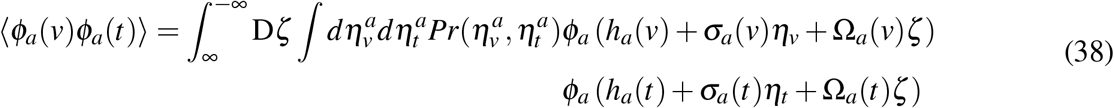

where

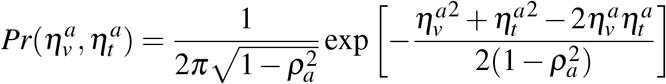

is a bivariate Gaussian probability density function with covariance

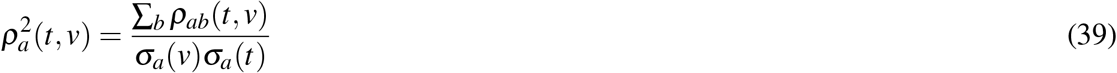

with *ρ*_*ab*_(*t, v*) given by Eq. 22.

We solved the DMFEs to generate Fig. S6.

### 4.2 Storage Capacity

Next, we estimate the storage capacity, the maximal number of sequences that a network can store. From the equations for the overlap dynamics (Eq. 28, 32, and 33), we see that if the factors 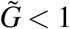, then the peak overlap decays with the pattern index. A retrieved sequence cannot be finished in that case, given enough sequence length. We further observe that

1. Effective learning strength *Ã* is insensitive to memory load for small *A*_0_ (Eq. 16).
2. Input variance *σ* ^2^ scales linearly with memory load (Eq. 23).
3. For *c* = *θ*_*a*_, *G*_*a*_(*c, σ* ^2^) decays monotonically with input variance for large input variance (Eq. 29). Otherwise, *G*_*a*_(*c, σ* ^2^) increases for small *σ* ^2^ and then decays with input variance.

Together, these observations indicate that there exists a maximal memory load such that in all conditions,

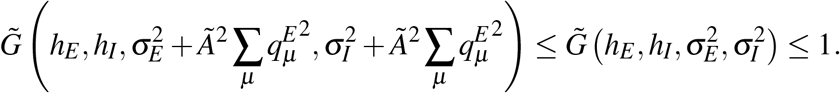

We can define the theoretical storage capacity 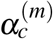 of a model *m* as

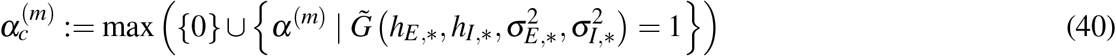

where *h*_∗_ and 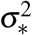 are the stationary input means and variances following Eq. 18 and Eq. 21, and the stationary means and second moments of firing rates satisfy the self-consistent equations

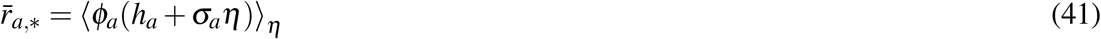

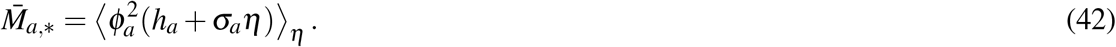

### 4.3 Network Simulations

#### 4.3.1 Initialization of the Network

For network simulations in Fig. 1, S1, S2, S3 and S5, neurons are initialized randomly, i.e. **r**^*a*^(0) = *φ*_*a*_(*η*), where *η* ∼ *N*(0, *a***I**). The initial pattern of the retrieved sequence *s* is presented as the external input to the excitatory population for the first 20 ms, i.e. 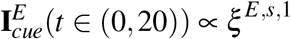. Initializing the inhibitory population with *ξ* ^*I*,*s*,1^ or presenting *ξ* ^*I*,*s*,1^ as the external input to the inhibitory population 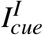 does not produce qualitatively different behaviors in the models considered in this study.

For robustness experiments, to ensure different models have the same maximal correlation with the first pattern (the same initial cue for retrieval), the excitatory neurons are initialized with 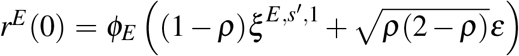, where *ρ* is the perturbation strength and *ε* is white noise. The first pattern is not presented later in **I**_*ext*_. The inhibitory neurons are initialized randomly as described above.

#### 4.3.2 Choices of Time Constants

Throughout this study, we assume that the membrane time constant for excitatory neurons, *τ*_*E*_ is much larger than that for inhibitory neurons, *τ*_*I*_, i.e. inhibitory neurons are much faster. The figures were generated with physiologically plausible values *τ*_*E*_ = 25 ms and *τ*_*I*_ = 5 ms [50]. Other choices (e.g. *τ*_*E*_ = 35 ms, *τ*_*I*_ = 10 ms) did not qualitatively alter the behaviors, except that the time scales are different.

#### 4.3.3 Choices of Connection Probability

Neurons in all networks are sparsely connected with connection probability 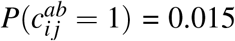. We chose this value because it allows simulations of the networks to be efficient, and the mean excitatory and inhibitory inputs to be large (given *N > O*(10^4^)). Other choices between 0.01 and 0.05 produce similar results.

#### 4.3.4 Choices of Transfer Functions

The transfer functions, *ϕ*_*E*_ and *ϕ*_*I*_, are sigmoidal functions and have the general form

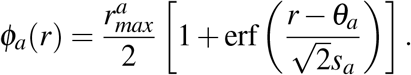

It is suggested that the parameters for the transfer functions for cortical excitatory and inhibitory neurons after normalizing inputs and outputs do not differ significantly [30]. Thus, we set *θ*_*a*_ = *θ* and *s*_*a*_ = *s* for all population *a*, but kept 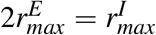 which matches qualitatively the maximal firing rates observed in cortex [30, 50]. *θ*, *s*, and 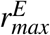 are adapted from [37] and is shown in Table 1.

#### 4.3.5 Approximation of the External Inputs

For efficiency, the external input for each neuron at each time step is drawn from an i.i.d. normal distribution with mean 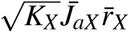 and standard deviation 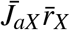. Generating the external inputs in another way, by first generating **r**^*X*^ and the connection matrices *J*_*aX*_, and then calculate the inputs *J*_*aX*_ **r**^*X*^, does not lead to qualitatively different dynamics.

#### 4.3.6 Numerical Method for Simulation

The networks are simulated using Euler method with time step 1 ms. Smaller time steps (0.5 ms and 0.1 ms) are also explored, but only slightly affect the transient (*t <* 30 ms) and do not lead to qualitatively different dynamics.

### 4.4 Grid Search

To investigate how inhibitory plasticity affects storage capacity, grid search is performed to optimize the capacity for three models: EE, EI-IE, and EE-EI-IE. In EE model, only E→ E synapses are used to store sequences using temporally asymmetric Hebbian rule. In EI-IE model, E →I and I →E synapses are used to store sequences. EE-EI-IE is a combination of the previous two models, where E →E, E →I, and I → E are used. In both EI-IE and EE-EI-IE model, E →I synapses are shaped by symmetric anti-Hebbian rule and I→ E by asymmetric Hebbian rule, making the inhibitory pathway from E→ I→ E effectively asymmetric Hebbian. The other combinations of plasticity rules that results in effectively asymmetric Hebbian inhibitory pathway should produce the same grid search outcome, as explained below.

To estimate the optimal capacity for the three models, we searched over 9 learning-related parameters: 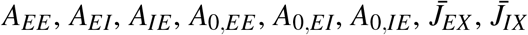 and 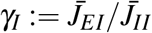 (Table 5). For a given set of parameters and each model, we gradually increased the memory load *α*^(*m*)^ (Eq. 9) by 0.01 per one

iteration, until 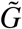 is both decreasing (smaller compared to previous iteration) and smaller than 1 (see Section 4.2). For each load value, the stationary means 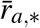 and second moments *M*_*a*,∗_ for the two populations are solved iteratively using Eq. 41 and 42.

To avoid missing solutions with even larger capacities, we did “finer” grid searches concentrated around the solutions with non-zero capacities found by the “coarser” search presented in Fig. S8. Specifically, *A*_0,*EE*_ ≤0 for the EE model, *A*_0,*EI*_, *A*_0,*IE*_≤ 0 for the EI-IE model, and *A*_0,*EE*_, *A*_0,*IE*_ ≤ 0, *A*_0,*EI*_ ≥0 for the EE-EI-IE model (Fig. S9). These finer grid searches did not lead to substantially different optimal capacities and capacity distributions.

### 4.5 Robustness

#### 4.5.1 Experiment Setup

A model could be very robust if its memory load is far below its capacity. It is more helpful to compare models with load close to their capacity. To do that, each model stores two sequences: one has a fixed length (12 patterns) and is retrieved, while the length of the other sequence is maximized and depends on the model types and parameters. With the guidance of theoretical capacities calculated in the previous sections, we increased the length of the second (un-retrieved) sequence until the maximal correlation with the last pattern of the retrieved sequence is slightly above 0.1 under small perturbation and noise.

The parameter sets are given in Table 7 and 8. Notice that we slightly adjusted their parameters such that the stationary mean excitatory rates are 5 ∼ 10 Hz, and the stationary mean inhibitory rates are 15 ∼ 30 Hz.

#### 4.5.2 Measuring the Robustness

To quantify robustness, we devised the following measures:

1. Maximal correlation between the excitatory neuron firing rates and the last pattern.
2. A binary variable whether the peaks of correlation curves are in the right order as the index of patterns.
3. Correlation peak definiteness, defined by 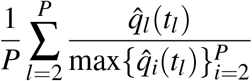 where 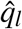 is the correlation with the *l*-th pattern and *t*_*l*_ = argmax_*t*_ 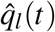 is the peak time. Ideally, when each correlation reaches its maximum at a particular time, it should also be larger than the correlation with the other patterns and this quantity is 1.
4. Average of maximal correlation with each pattern.
5. Resolution, defined as 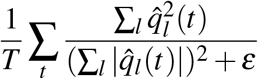 where *ε* ≈ 0.

### 4.6 Chaos

Previous studies on balanced E-I networks with random weights show that the strong synaptic weights necessary for the global E-I balance result in chaotic dynamics [51]. We check whether our networks also have chaotic dynamics via simulations. Each model (EE, EI-IE, or EE-EI-IE) has the same network size (*N*_*E*_ = *N*_*X*_ = 40000 and *N*_*I*_ = 10000), same connectivity probability (*c* = 0.015), same constant external inputs 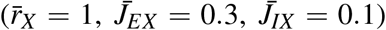, and same relative memory load 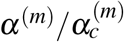. For each generated network, we simulated it with an unperturbed and a perturbed version of the first pattern of a stored sequence. That is, the initial conditions are:

1. Unperturbed: 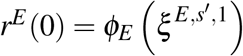
2. Perturbed: 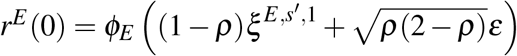

where we set the perturbation strength to be small, *ρ* = 0.001, and simulated for sufficient long time. There is no other source of stochasticity. To quantify the dynamics, we calculated the Euclidean distance, 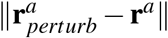, and cosine similarity, 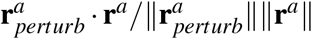, between the population vectors from the perturbed and unperturbed cases, where ‘·’ denotes dot product and *a* ∈ {*E, I*}.

The simulations show that small difference in the initialization is magnified over time (Fig. S7), indicating that the networks have chaotic dynamics.

## 5 Supplementary Figures

**Figure S1:**
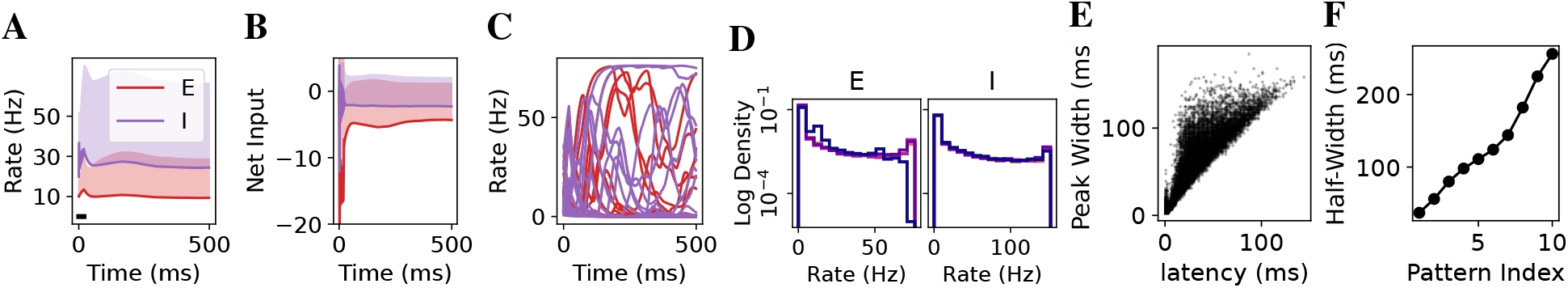
Characteristics of the EE model. (A) The mean rates (solid lines) of excitatory (red) and inhibitory (purple) neurons over time. The vertical width of the shaded area marks 1 standard deviation of firing rates. (B) The means and standard deviations of inputs to excitatory and inhibitory neurons. (C) Examples of excitatory and inhibitory neuron firing rates. (D) Distributions of excitatory (left) and inhibitory (right) firing rates. From dark to bright, the distributions at *t* = 50, 150, 250, 350, 450 ms. (E) The widths of peaks in the excitatory firing rates are plotted against the latencies of the corresponding peaks.(F) The half-width of the correlation curves as a function of pattern index. Half-width is defined by the width at the half of maximal correlation for each pattern. Parameters in Table 3.

**Figure S2:**
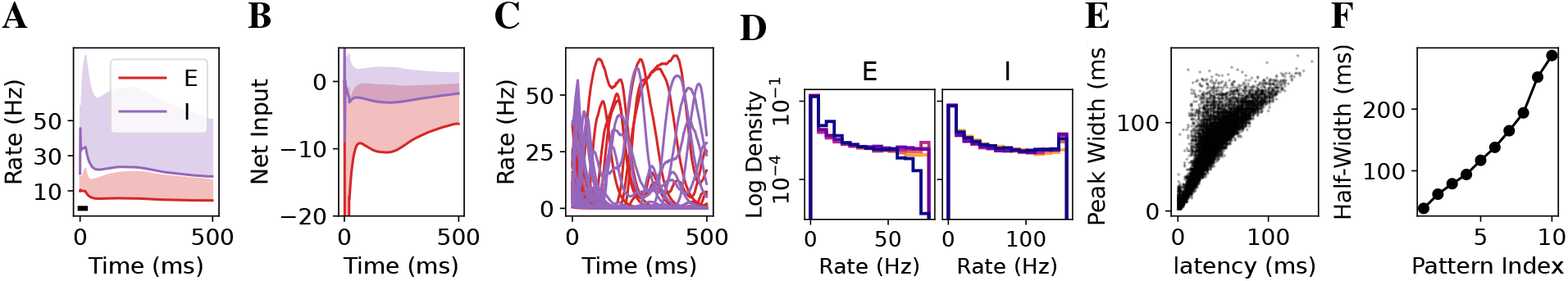
Characteristics of the EI-IE model. The notations of colors and lines are the same as Fig. S1. Parameters in Table 3.

**Figure S3:**
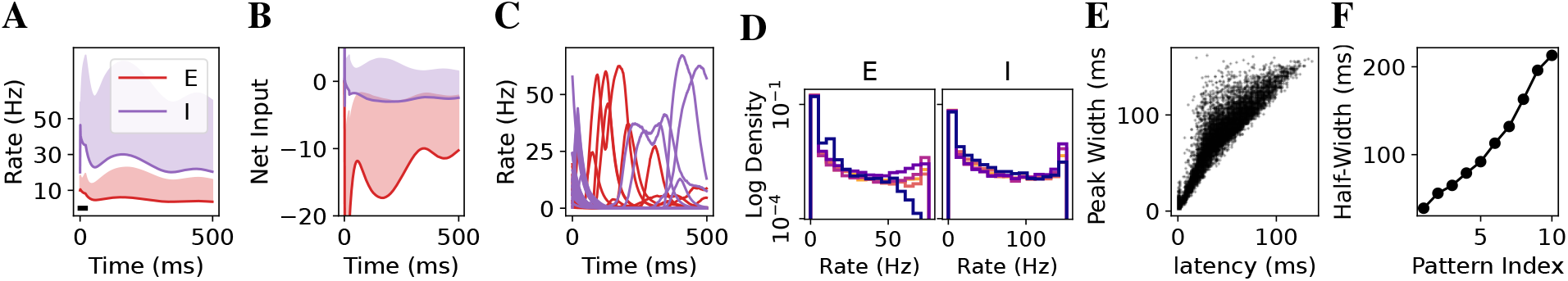
Characteristics of the EE-EI-IE model. The notations of colors and lines are the same as Fig. S1. Parameters in Table 3.

**Figure S4:**
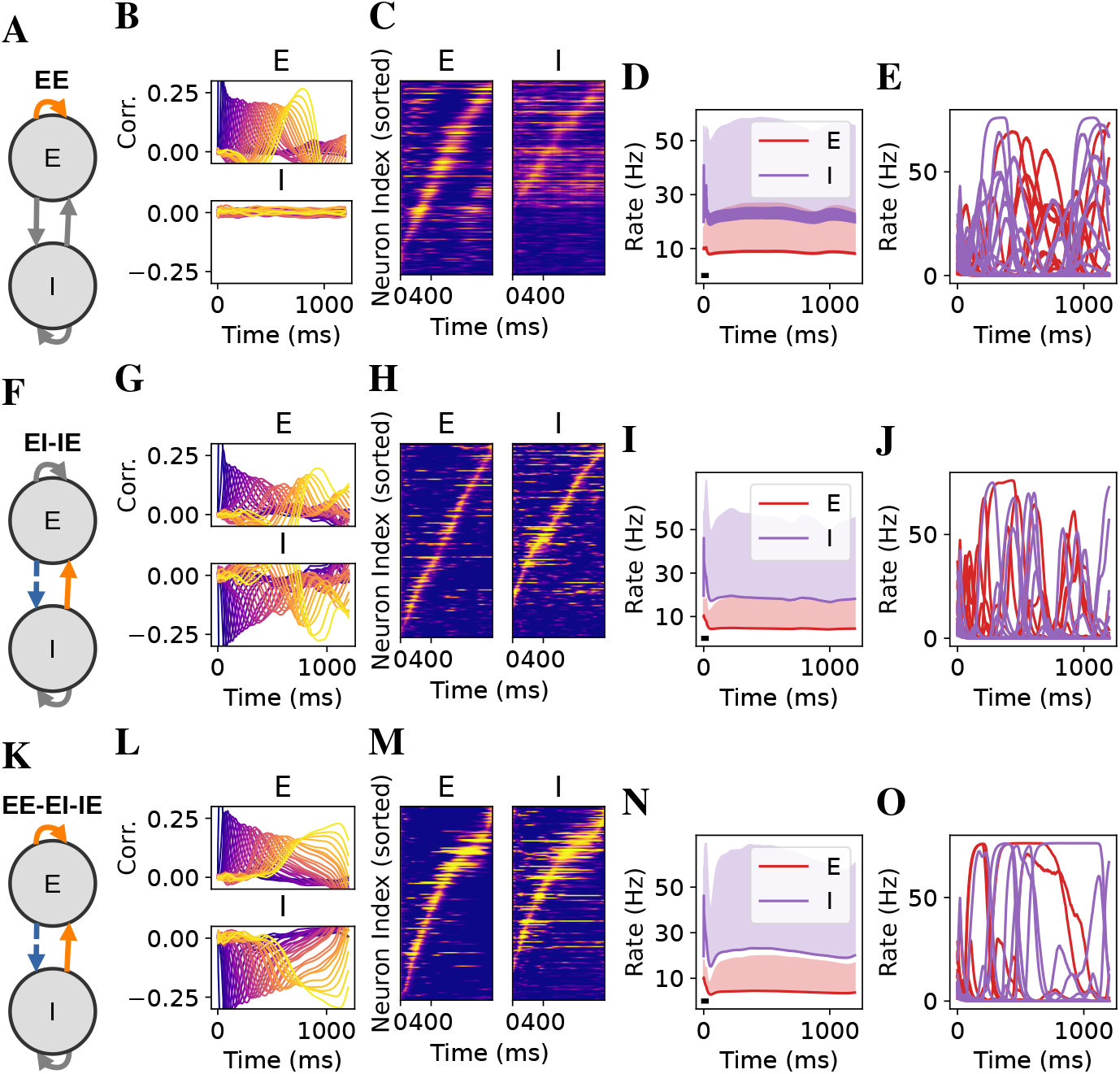
Models can store and retireve much longer sequences (*P* = 30). From left to right, the schematic of the network, correlations with sequence patterns, raster plot of active neurons, the means and standard deviations of firing rates, and examples of excitatory and inhibitory neuron firing rates for (A-E) EE model, (F-J) EI-IE model, and (K-O) EE-EI-IE model. Parameters in Table 4.

**Figure S5:**
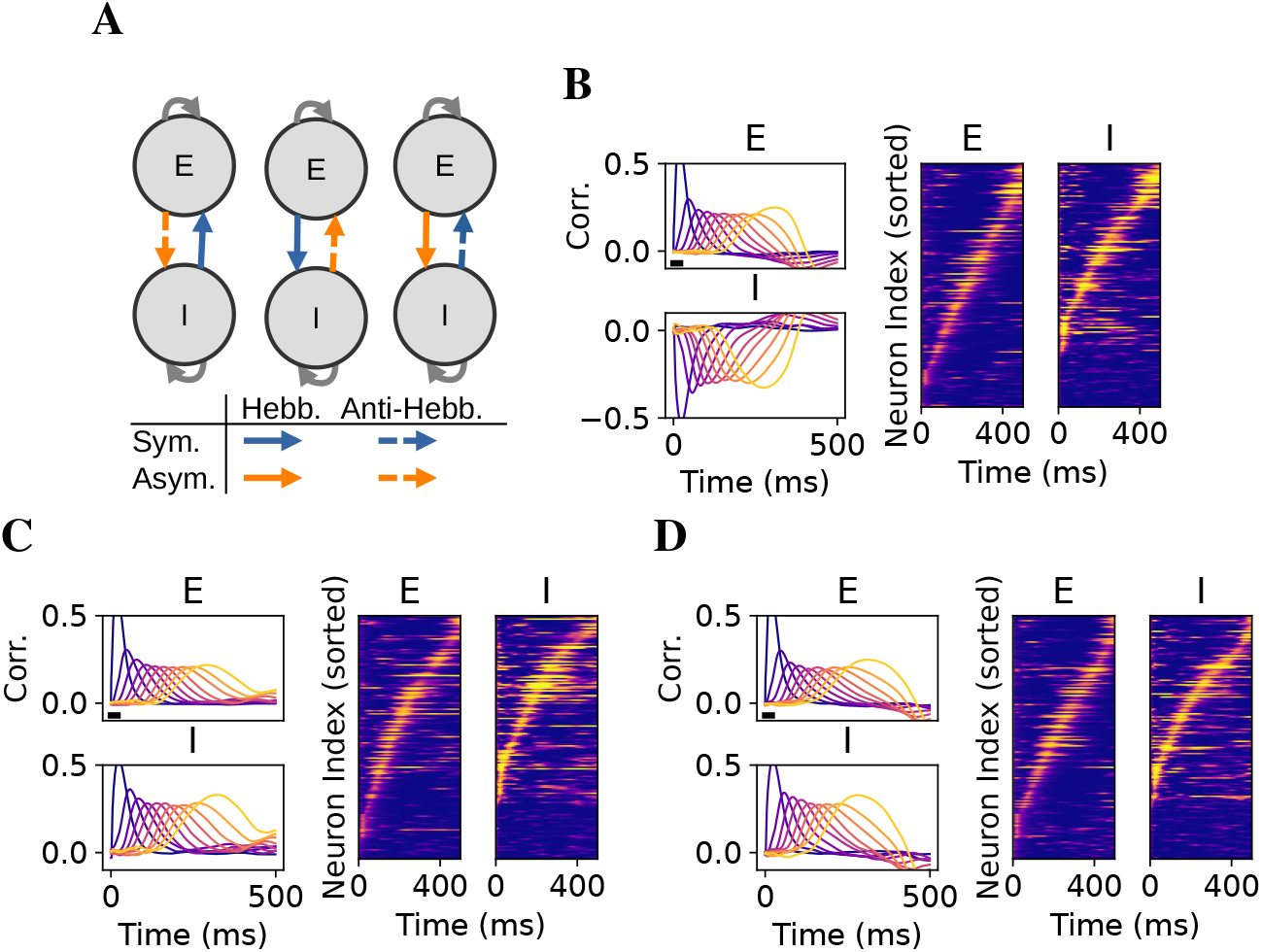
Different combinations of plasticity rules lead to similar retrieval dynamics in the excitatory neurons as long as the pathways are effectively asymmetric Hebbian. (A) Schematics of different combinations of plasticity rules that lead to effectively asymmetric Hebbian. The schematics from left to right correspond to (B), (C), and (D), respectively. (B-D) The correlations with the patterns in the retrieved sequence (left) and the raster plots (right). Parameters are the same as the EI-IE model in Fig. S2.

**Figure S6:**
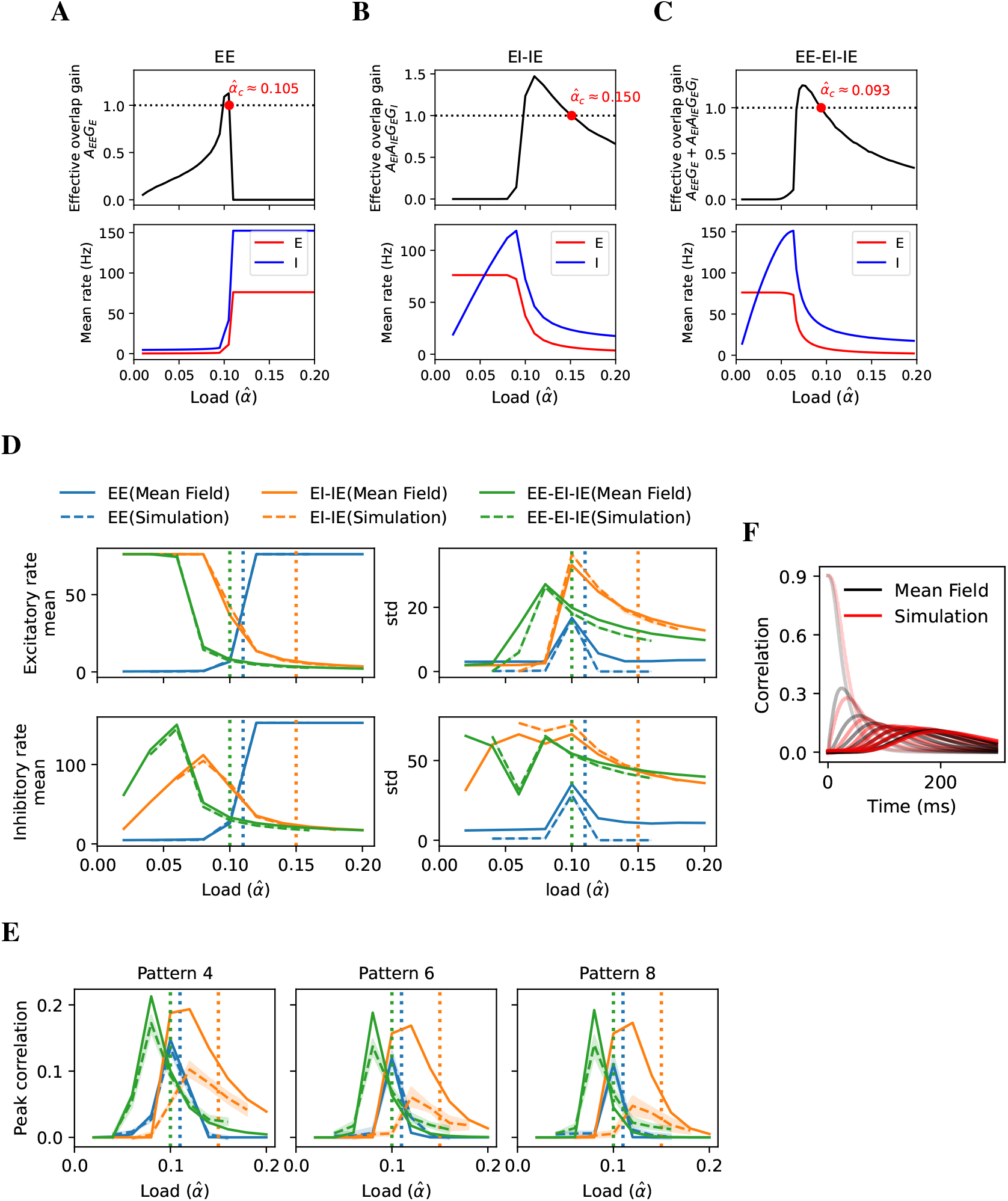
Estimation of storage capacity using the method in Section 4.2, simulations, and DMFEs. (A-C) The theoretical storage capacity (Section 4.2) of the EE (A), EI-IE (B), and EE-EI-IE (C) models. Red dots mark the capacity values, which are close to the optima. (D) The means and standard deviations of excitatory and inhibitory firing rates solved using the DMFEs (solid lines) are qualitatively similar to those from the simulations (dashed lines) of actual networks for different memory load. Dotted lines mark the theoretical storage capacity (Section 4.2). (E) The maximal correlations with different patterns also matches, especially for large load values. Importantly, when the memory load is larger than the theoretical storage capacity for each model (Section 4.2), the maximal correlations for later patterns drop sharply to zero. (F) An example of the retrieval dynamics from DMFE (black) and simulation (red) of the EE model at *α*^(*m*)^ = 0.1. Parameters are in Table 6.

**Figure S7:**
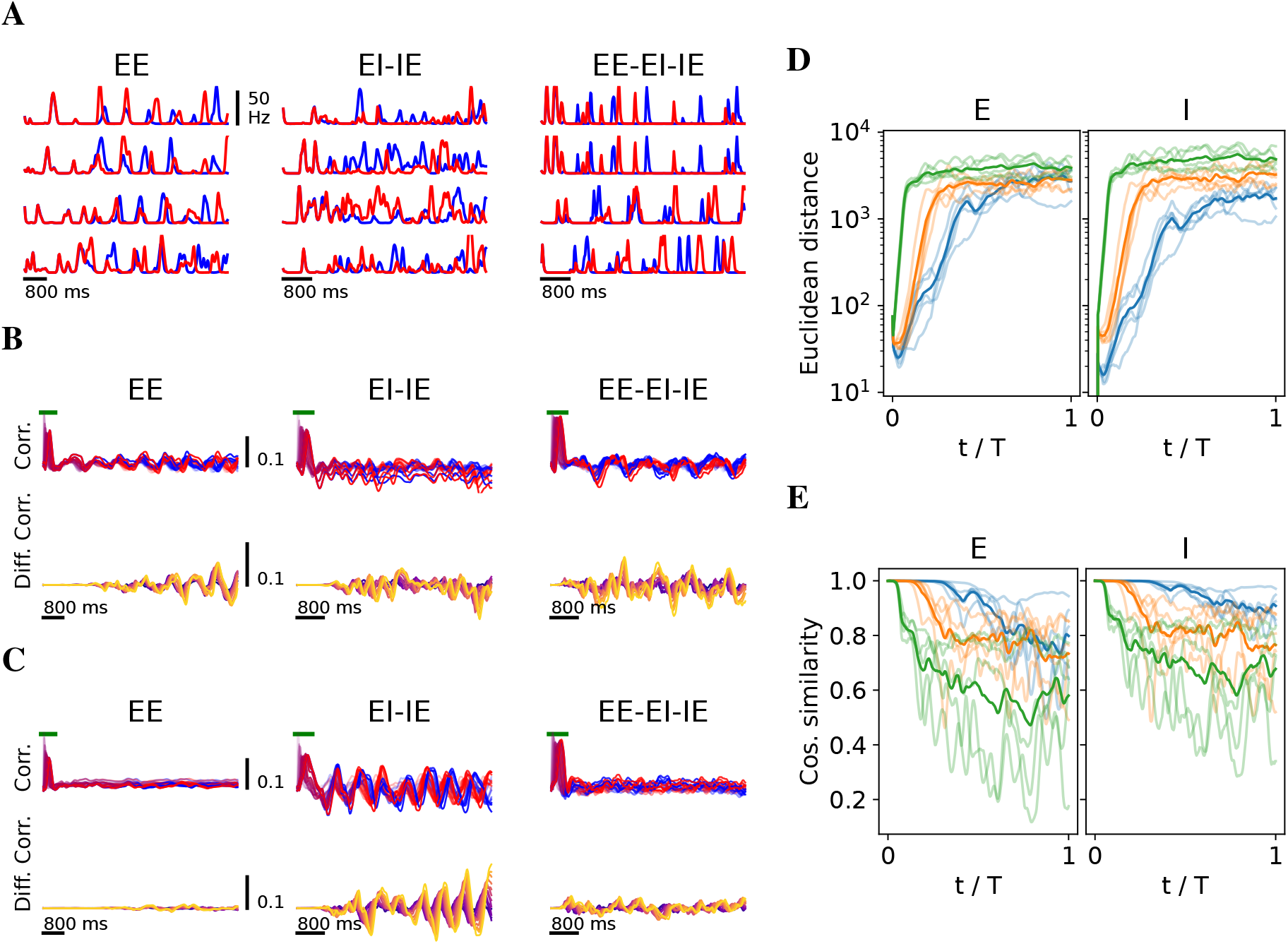
Chaotic behaviors in the networks. (A) Firing rates of 4 randomly chosen neurons in each model when the network is initialized to the first pattern of a sequence (blue) or a slightly perturbed version of it (red; perturbation strength *ρ* = 0.001). (B) Top row, the correlations between the excitatory neuron firing rates and the patterns of the retrieved sequence in the unperturbed (blue) and perturbed (red) cases. Green bar indicates sequence retrieval. Bottom row, the differences in the correlations between the unperturbed and perturbed cases. Brighter curve corresponds to later pattern in the sequence. (C) As (B), but for another set of randomly generated patterns and connectivity. (D) The Euclidean distance between the population vectors from the perturbed and unperturbed cases, i.e. 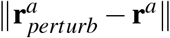. (E) The cosine similarity between the population vectors from the perturbed and unperturbed cases, i.e.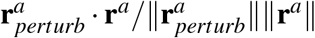. Parameters are in Table 7, though other parameters lead to similar chaotic dynamics.

**Figure S8:**
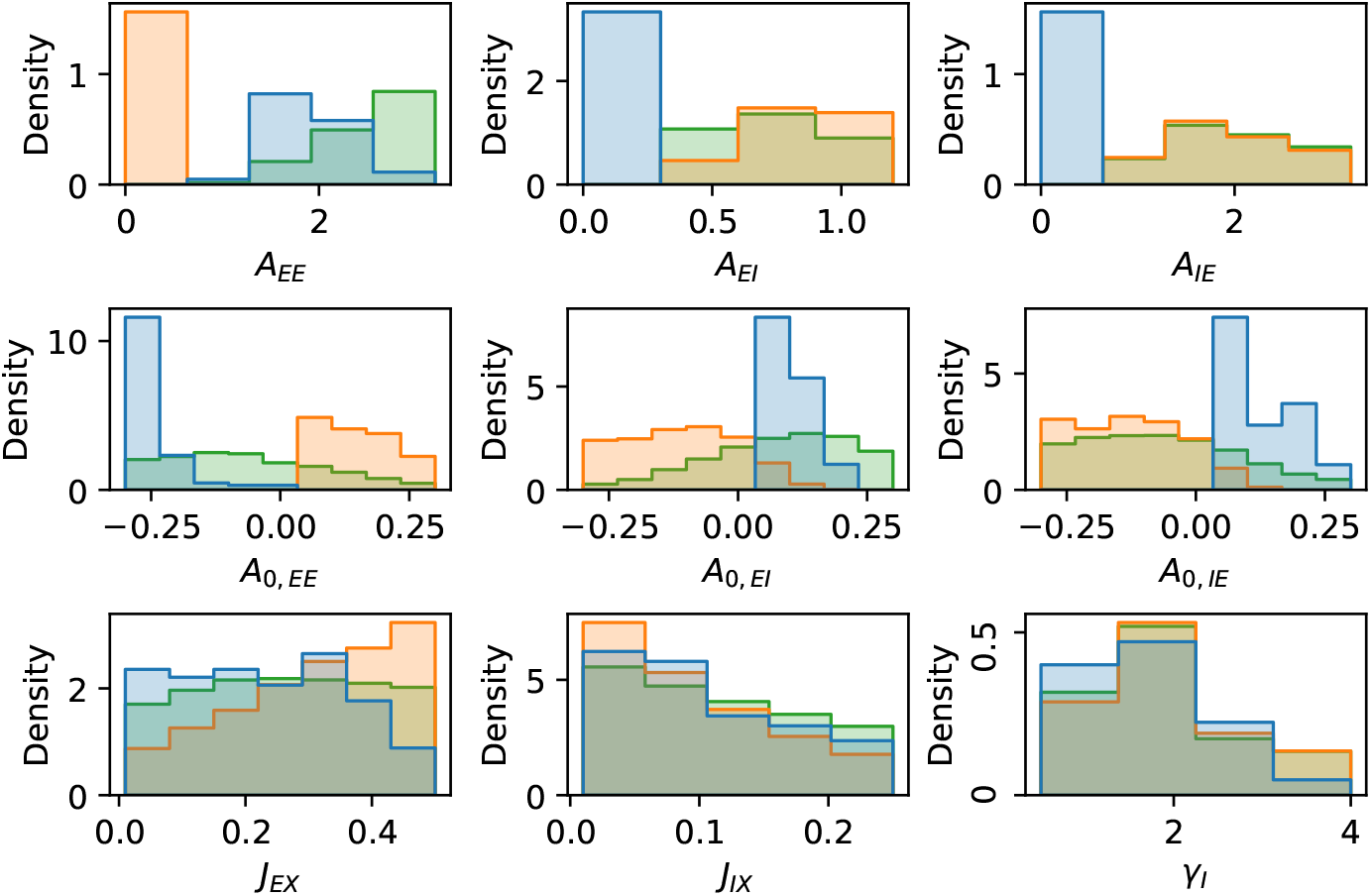
For each model, the distributions of parameters for good solutions whose capacities are above 80% of the model’s optimal capacity. Blue, EE model; orange, EI-IE model; green, EE-EI-IE model.

**Figure S9:**
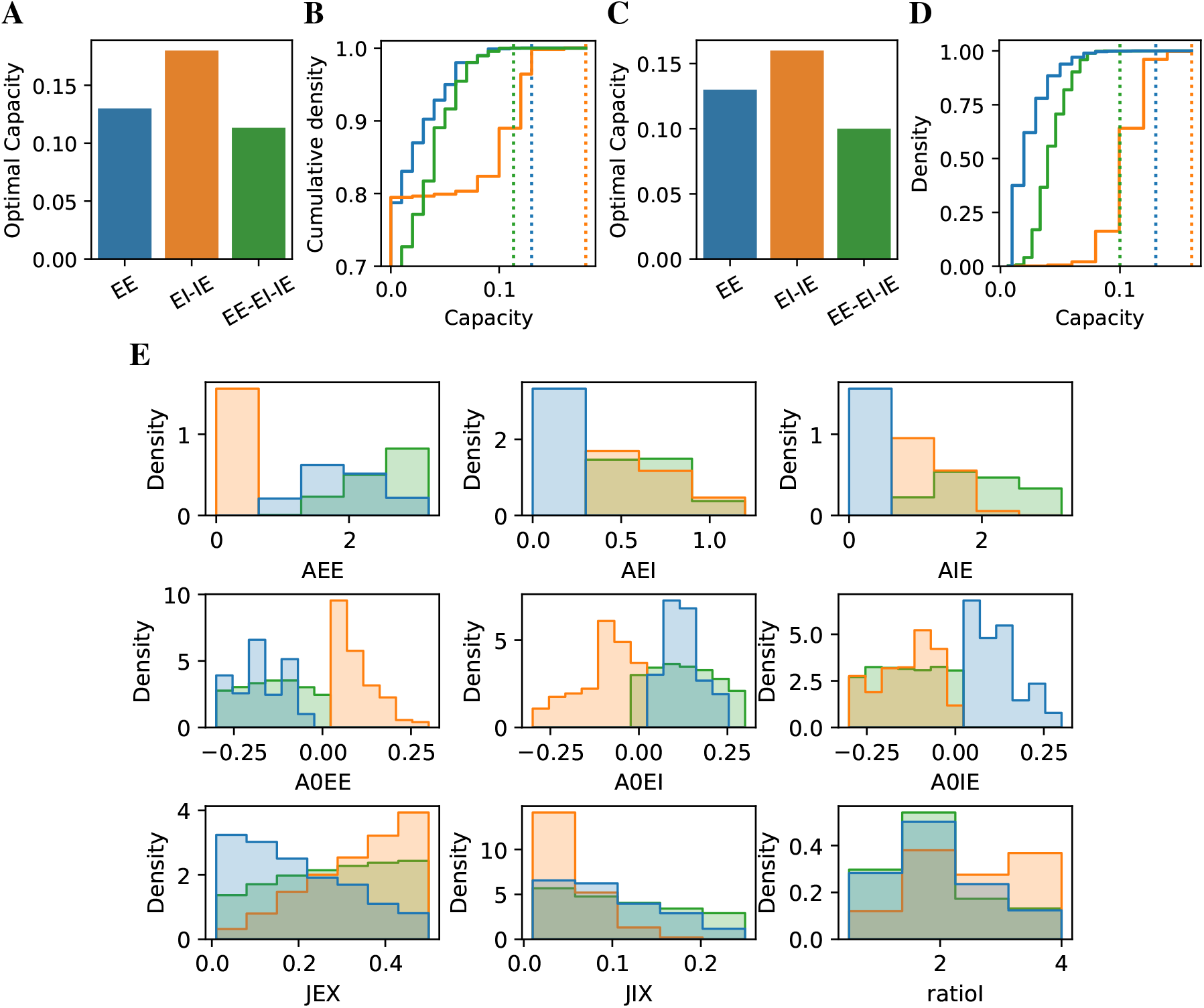
The results from a finer grid search where the ranges of *A*_0_’s are model-specific, and chosen based on the results of the grid search in Fig. 2. For EE model, *A*_0,*EE*_∈ [ −0.3, 0], *A*_0,*EI*_, *A*_0,*IE*_∈ [0, 0.3]. For EI-IE model, *A*_0,*EE*_∈ [0, 0.3], *A*_0,*EI*_, *A*_0,*IE*_∈ [ −0.3, 0]. For EE-EI-IE model, *A*_0,*EE*_, *A*_0,*IE*_∈ [ −0.3, 0], *A*_0,*EI*_ ∈ [0, 0.3]. (A) The optimal capacity for each model (B) The cumulative distribution of capacities for each model across all valid searched parameter sets. (C-D) as (A-B) but the parameter sets that can lead to unrealistic firing rates are excluded. We consider 1 ∼10 Hz as realistic excitatory firing rate, and 10∼30 Hz as realistic inhibitory firing rate. (E) For each model, the distributions of parameters for good solutions whose capacities are above 80% of the model’s optimal capacity.

**Figure S10:**
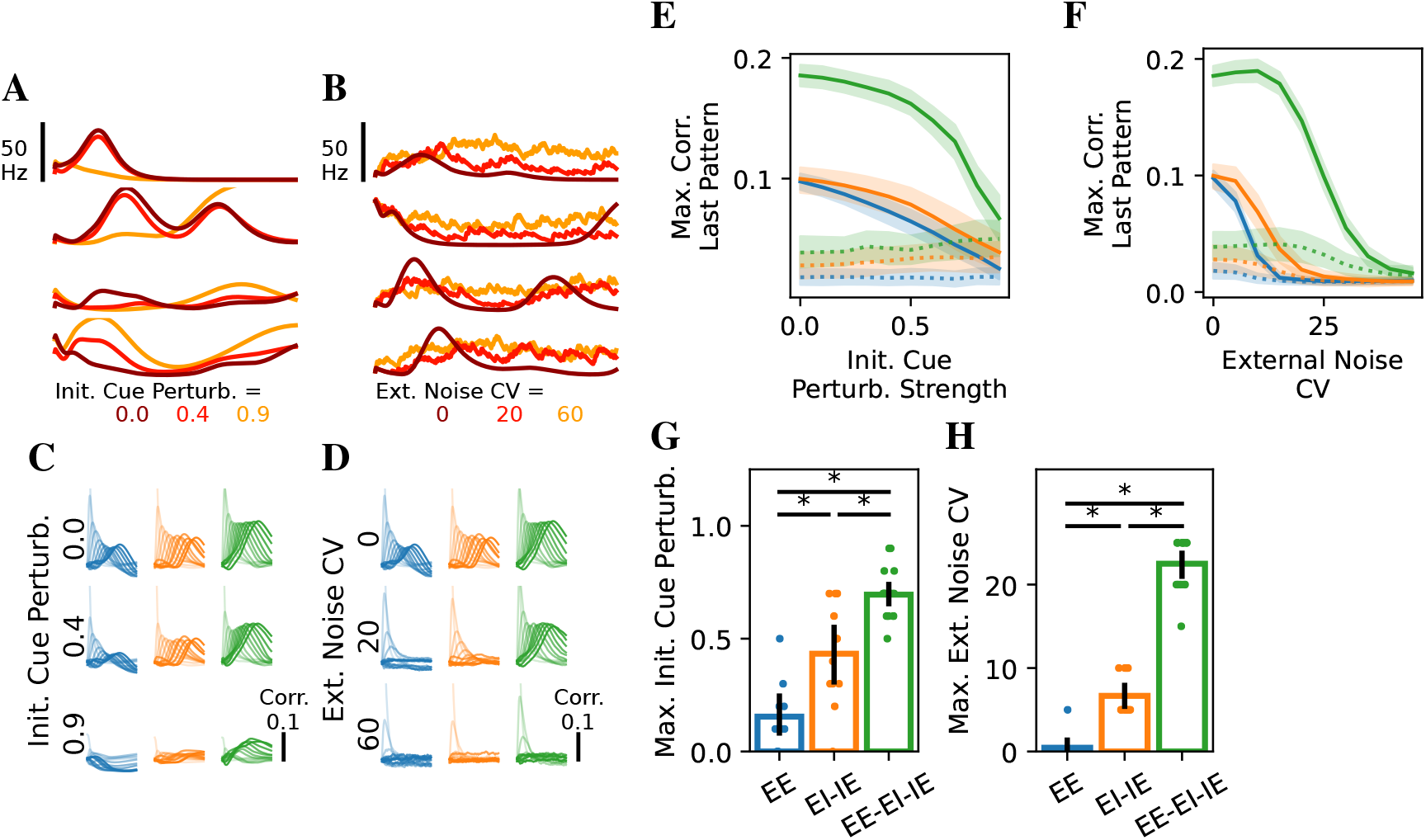
Robustness to initial cue perturbation. As Fig. 4, but for another parameter set (Table 8).

**Figure S11:**
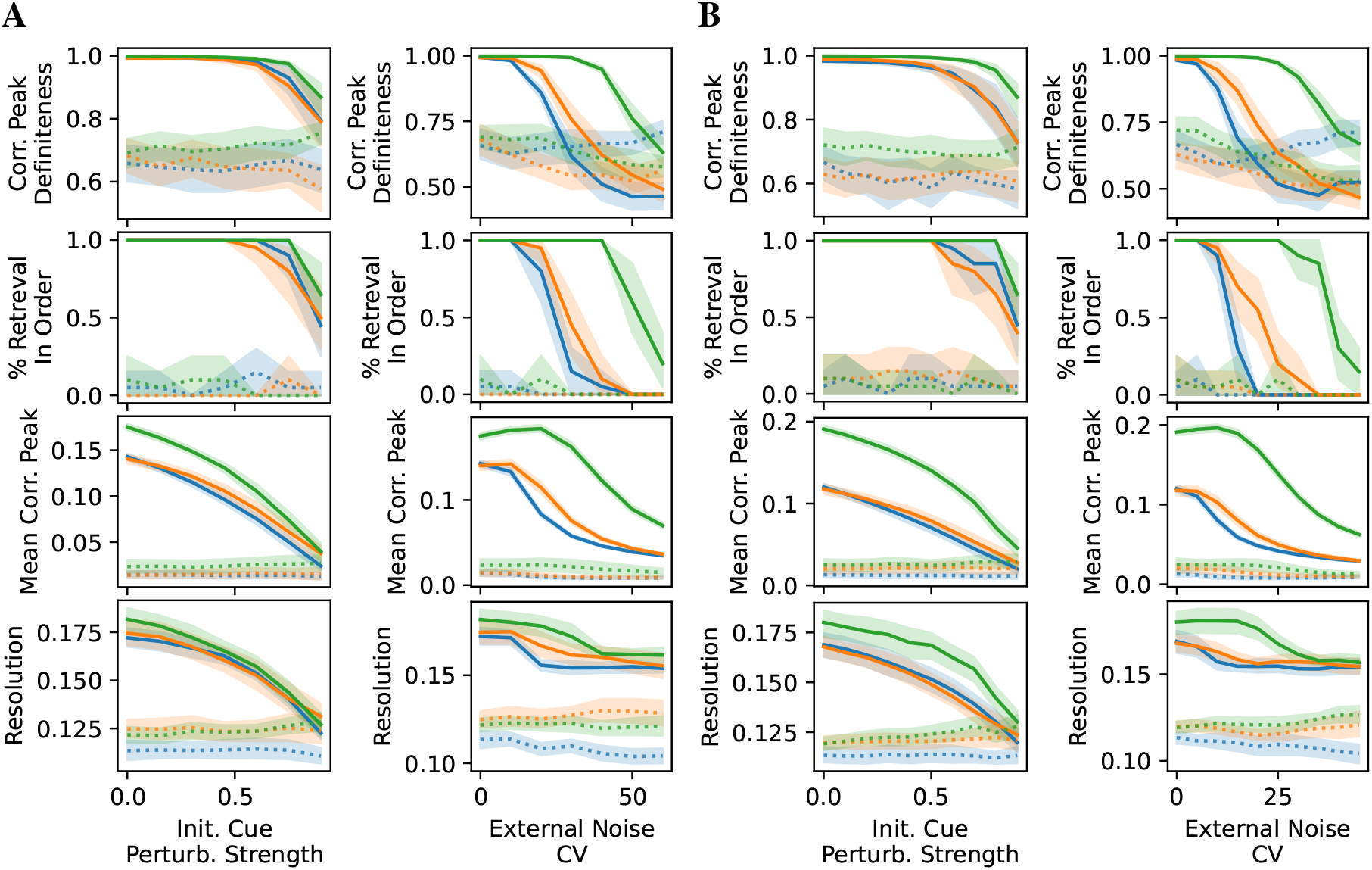
Additional measurements for robustness experiments in (A) Fig. 4 and (B) Fig. S10.

## 6 Parameter Tables

**Table 1:** Parameters for the transfer functions used throughout the study 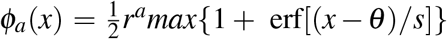.

| Symbol | Value | Description |
| --- | --- | --- |
| $s$ | 1.946 | Inverse slope at the mid point |
| $\theta$ | 2.46 | Mid point offset |
| $r_{max}^E$ | 76.2 | Maximal firing rate for excitatory neurons |
| $r_{max}^I$ | 152.4 | Maximal firing rate for inhibitory neurons |

**Table 2:**
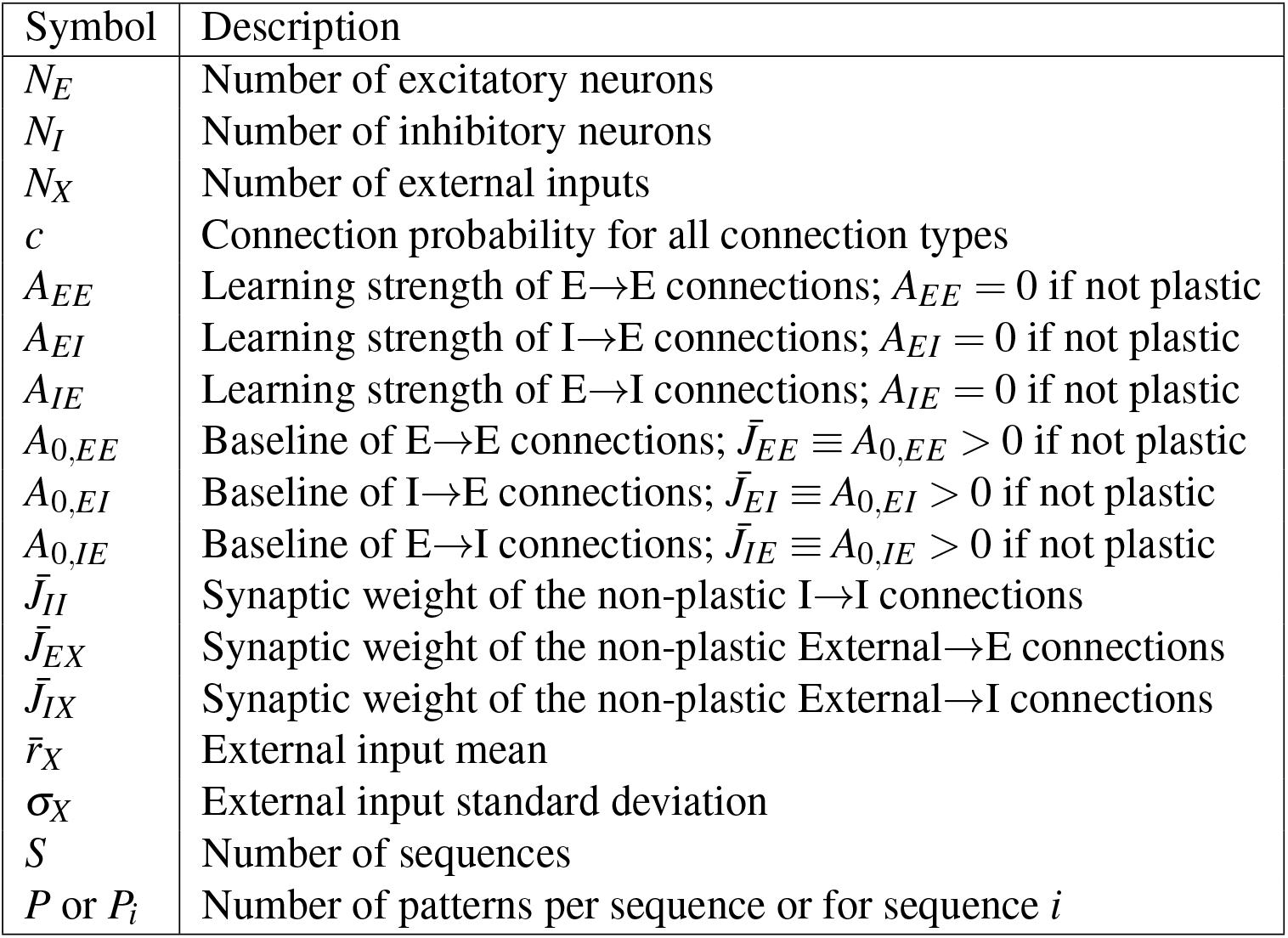
Explanation of all parameters.

**Table 3:**
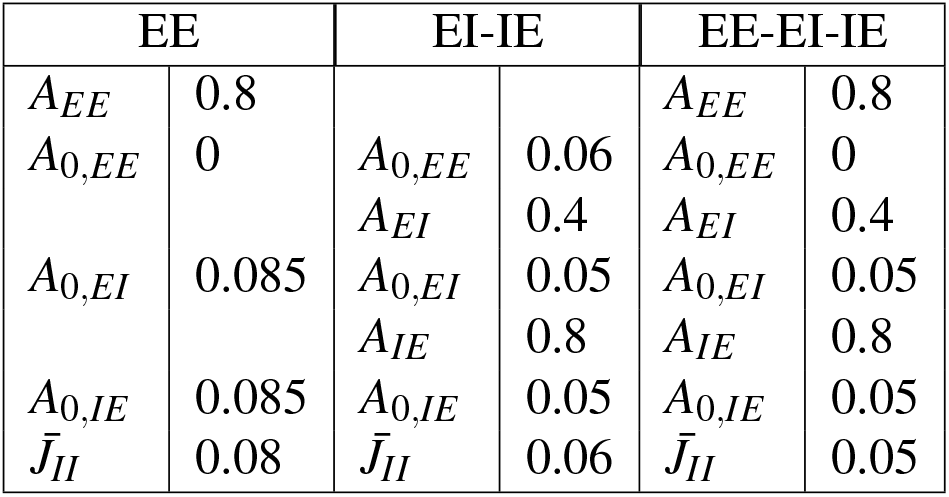
Parameters for the models in Fig. 1, S1, S2, S5, and S3. Other parameters are the same across all models: *N*_*E*_ = *N*_*X*_ = 40000, *N*_*I*_ = 10000, *c* = 0.015, *S* = 2, *P* = 10, 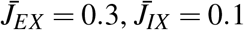, 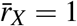, and *σ*_*X*_ = 1.

**Table 4:**
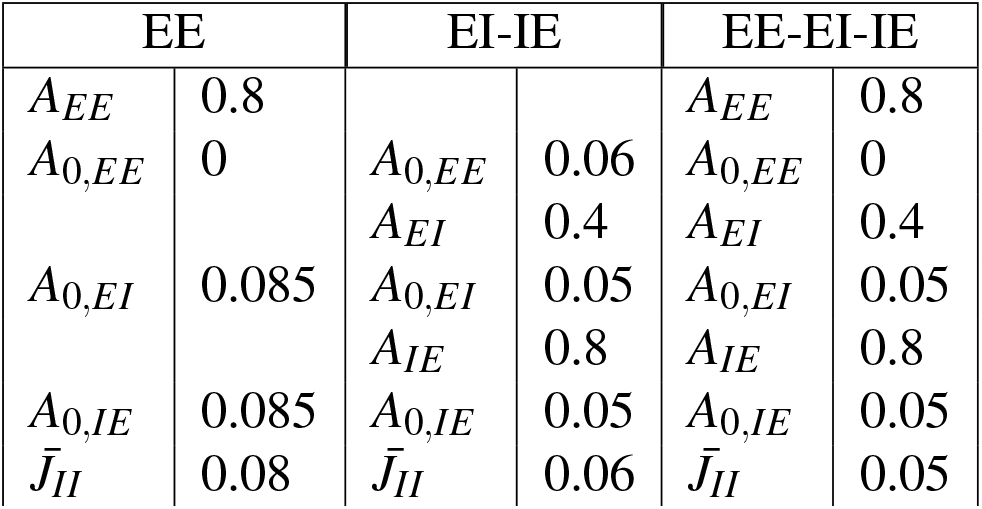
Parameters for the models storing long sequences in Fig. S4. Other parameters are the same across all models: *N*_*E*_ = *N*_*X*_ = 40000, *N*_*I*_ = 10000, *c* = 0.03, *S* = 2, *P* = 30, 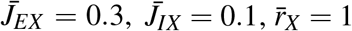, and *σ*_*X*_ = 1.

**Table 5:**
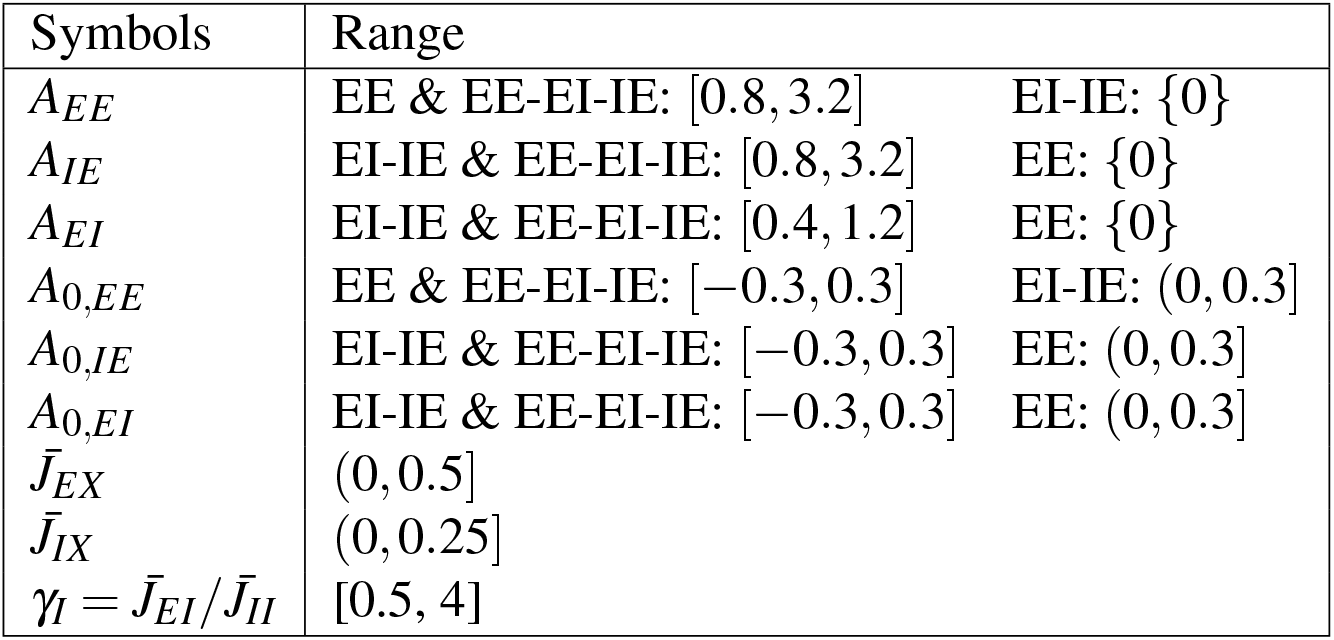
Parameters in grid search and their ranges.

**Table 6:** Parameters for Fig. S6. Other parameters are the same across all models: *N*_*E*_ = *N*_*X*_ = 40000, *N*_*I*_ = 10000, *c* = 0.04, 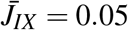and 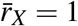.

| EE |  | EI-IE |  | EE-EI-IE |  |
| --- | --- | --- | --- | --- | --- |
| $A_{EE}$ | 1.6 | | | $A_{EE}$ | 1.6 |
| $A_{0,EE}$ | -0.22 | $A_{0,EE}$ | 0.09 | $A_{0,EE}$ | -0.05 |
| | | $A_{EI}$ | 0.6 | $A_{EI}$ | 0.6 |
| $A_{0,EI}$ | 0.075 | $A_{0,EI}$ | 0 | $A_{0,EI}$ | 0 |
| | | $A_{IE}$ | 1.6 | $A_{IE}$ | 1.6 |
| $A_{0,IE}$ | 0.075 | $A_{0,IE}$ | -0.22 | $A_{0,IE}$ | -0.22 |
| $\bar{J}_{II}$ | 0.04 | $\bar{J}_{II}$ | 0.06 | $\bar{J}_{II}$ | 0.085 |
| $\bar{J}_{EX}$ | 0.1 | $\bar{J}_{EX}$ | 0.6 | $\bar{J}_{EX}$ | 0.6 |

**Table 7:**
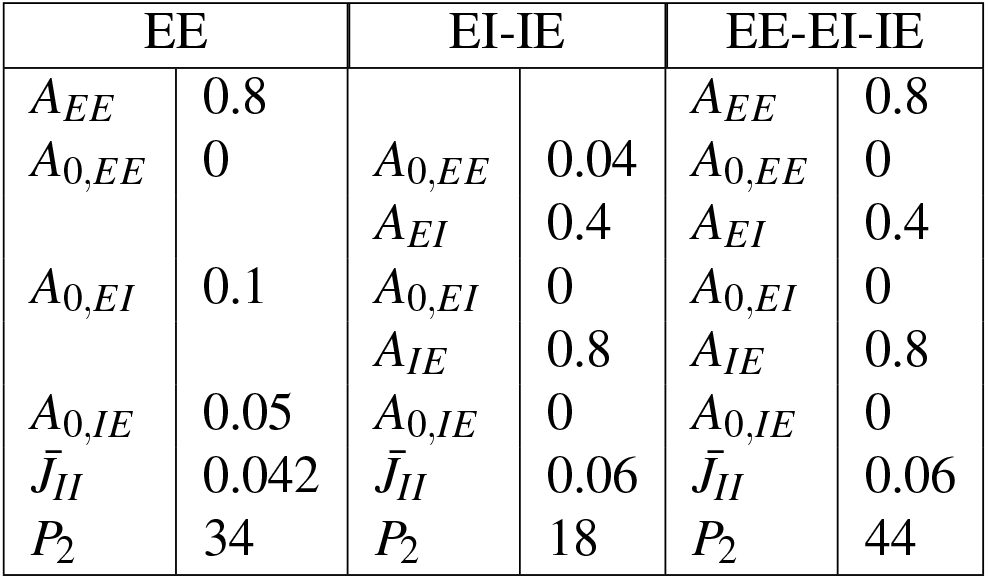
Parameters for the models in robustness comparisons in Fig. 4. Other parameters are the same across all models: *N*_*E*_ = *N*_*X*_ = 40000, *N*_*I*_ = 10000, *c* = 0.015, 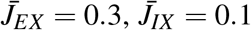 and 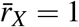. The theoretical capacities are approximately 0.1 (EE), 0.13 (EI-IE), and 0.08 (EE-EI-IE).

**Table 8:** Parameters for the models in robustness comparisons in Fig. S10. Other parameters are the same across all models: *N*_*E*_ = *N*_*X*_ = 40000, *N*_*I*_ = 10000, *c* = 0.015, 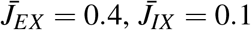 and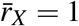. The theoretical capacities are approximately 0.1 (EE), 0.15 (EI-IE), and 0.09 (EE-EI-IE).

| EE |  | EI-IE |  | EE-EI-IE |  |
| --- | --- | --- | --- | --- | --- |
| $A_{EE}$ | 0.8 | | | $A_{EE}$ | 0.8 |
| $A_{0,EE}$ | -0.06 | $A_{0,EE}$ | 0.03 | $A_{0,EE}$ | -0.06 |
| | | $A_{EI}$ | 0.4 | $A_{EI}$ | 0.4 |
| $A_{0,EI}$ | 0.06 | $A_{0,EI}$ | -0.03 | $A_{0,EI}$ | -0.03 |
| | | $A_{IE}$ | 0.8 | $A_{IE}$ | 0.8 |
| $A_{0,IE}$ | 0.07 | $A_{0,IE}$ | -0.06 | $A_{0,IE}$ | -0.06 |
| $\bar{J}_{II}$ | 0.042 | $\bar{J}_{II}$ | 0.03 | $\bar{J}_{II}$ | 0.045 |
| $P_2$ | 26 | $P_2$ | 17 | $P_2$ | 41 |

## Footnotes

1 For non-plastic connections *J*^*ab*^, *A*_*ab*_ = 0 and 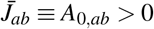

## Notes

### Competing Interest Statement

The authors have declared no competing interest.

